# Proteome-wide cellular thermal shift assay reveals novel crosstalk between brassinosteroid and auxin signaling

**DOI:** 10.1101/2021.12.14.472597

**Authors:** Qing Lu, Yonghong Zhang, Joakim Hellner, Xiangyu Xu, Jarne Pauwels, Caterina Giannini, Qian Ma, Wim Dejonghe, Huibin Han, Brigitte Van de Cotte, Francis Impens, Kris Gevaert, Jiří Friml, Ive De Smet, Daniel Martinez Molina, Eugenia Russinova

## Abstract

Despite the growing interest in using chemical genetics in plant research, small-molecule target identification remains a major challenge. The cellular thermal shift assay coupled with high-resolution mass-spectrometry (CETSA MS) that monitors changes in the thermal stability of proteins caused by their interactions with small molecules, other proteins, or post-translational modifications allows the identification of drug targets, or the study of protein-metabolite and protein-protein interactions mainly in mammalian cells. To showcase the applicability of this method in plants, we applied CETSA MS to intact *Arabidopsis thaliana* cells and identified the thermal proteome of the plant-specific glycogen synthase kinase 3 (GSK3) inhibitor, bikinin. A comparison between the thermal- and the phospho-proteomes of bikinin revealed the auxin efflux carrier PIN-FORMED1 (PIN1) as a novel substrate of the *Arabidopsis* GSK3s that negatively regulate the brassinosteroid signaling. We established that PIN1 phosphorylation by the GSK3s is essential for maintaining its intracellular polarity that is required for auxin-mediated regulation of vascular patterning in the leaf thus, revealing a novel crosstalk between brassinosteroid and auxin signaling.

**Significance Statement:** Chemical genetics, which investigates the biological processes using small molecules, is gaining interest in plant research. However, a major challenge is to uncover the mode of action of the small molecule. Here, we applied the cellular thermal shift assay coupled with mass spectrometry (CETSA MS) to intact *Arabidopsis* cells and showed that bikinin, the plant-specific glycogen synthase kinase 3 (GSK3) inhibitor, changed the thermal stability of some of its direct targets and putative GSK3 interacting proteins. In combination with phosphoproteomics, we also revealed that GSK3s phosphorylate the auxin carrier PIN-FORMED1 (PIN1) and regulated its polarity that is required for the vascular patterning in the leaf.

## Introduction

Although the application of chemical genetics to plant research is gaining interest (1-3), unravelling the mode of action of the small molecules remains a major challenge. The cellular thermal shift assay (CETSA) is a label-free method that can assess target engagement directly in live cells (4), but its application to plant cells remains limited (5, 6). The technique is based on the biophysical principle that a ligand can induce changes in the thermal stability of the target protein, allowing the generation of so-called protein melting curves (7, 8). Similarly to the classical thermal shift assay with purified proteins, small-molecule binding typically leads to protein stabilization and an increase in melting temperature (T_m_). Coupling CETSA with multiplexed quantitative mass spectrometry (MS) enables the monitoring of an entire proteome for changes in protein thermostability in the presence of a small molecule (9, 10). Consequently, proteins interacting with this molecule can be identified without previous knowledge of the pathways or molecular mechanisms involved (11). As the thermal stability of a protein may also be affected by post-translational modifications or by binding to other proteins, cofactors, or metabolites, CETSA MS carried out on intact cells, in which active signaling takes place, allows the identification of effector proteins downstream of the direct target (9).

The small molecule bikinin (12) is an inhibitor specifically targeting the *Arabidopsis thaliana* Shaggy/ glycogen synthase kinase 3 (GSK3)-like kinases (*At*SKs), including the key negative brassinosteroid (BR) signaling regulator BR-INSENSITIVE2 (BIN2)/*At*SK21 that phosphorylates and inactivates two main transcription factors, BRASSINAZOLE RESISTANT1 (BZR1) and BRI1-EMS-SUPPRESSOR1 (BES1)/BZR2 (13, 14). The *Arabidopsis* genome encodes 10 *At*SKs, which belong to four groups (15) evidence exists for at least six, *At*SK11, *At*SK12, *At*SK13, *At*SK22, *At*SK23, and *At*SK32, that they negatively regulate the BR signaling just as BIN2/*At*SK21 (16–18). Likewise the mammalian GSK3s (19), *At*SKs also phosphorylate numerous substrates and control many developmental and physiological processes in plants such as root, stomatal and flower development, xylem differentiation, responses to light, and different abiotic and biotic stresses (15). BIN2/*At*SK21 also mediates the crosstalk between BR and other plant hormones, including auxin (15).

In spite of the reported interdependency and cooperation (20-25), the molecular mechanisms of the signaling crosstalk between BRs and auxin are still not well understood. Although auxin does not affect the phosphorylation state of the BZR1 and BES1/BZR2 transcription factors (14), BIN2/*At*SK21 interacts and phosphorylates the AUXIN RESPONSE FACTOR2 (ARF2) (21). This phosphorylation results in a loss in the DNA binding and repressor activity of ARF2 and facilitates auxin responses (21). BIN2/*At*SK21 also phosphorylates and activates ARF7 and ARF19 to promote lateral root development through an increase in auxin response (22). Moreover, BRs have been shown to control posttranscriptionally the endocytic sorting of PIN-FORMED 2 (PIN2) (23) and stimulate the nuclear abundance and signaling of auxin via repressing the accumulation of PIN-LIKES (PILS) proteins at the endoplasmic reticulum (24).

Here, by adapting the CETSA MS to *Arabidopsis* intact cells and combining it with the phosphoproteomics, we discovered that the auxin efflux carrier PIN-FORMED1 (PIN1) is a novel substrate of the *At*SKs. We found that phosphorylation mediated by the *At*SKs is required for PIN1 polarity and for leaf venation. In summary, we demonstrate that CETSA MS is a powerful method for identification of small-molecule targets as well as for discovery of new protein-protein interactions in plant cells.

## Results

### CETSA monitoring of small molecule-protein interactions in intact *Arabidopsis* cells

Previously, we had used the Western blot-based CETSA for small-molecule target validation in cell lysates of *Arabidopsis* seedlings (6). To extend the use of CETSA to intact plant cells, we adapted the available protocol (10) to *Arabidopsis* cell suspension cultures (Fig. S1). First, we tested whether the plant cell wall complicated the protein isolation by evaluating the efficiency of the freeze-thaw lysis method applied for mammalian cells (10). After washing and resuspension in protein extraction buffer, 100-µl aliquots of cells were freeze-thawed multiple times and the protein concentration of the supernatant was measured with the Bradford protein assay. The protein concentration in the lysate plateaued at the seventh freeze-thaw cycle (Fig. S2*A*). Moreover, considering that plants grow over a wider temperature range, we assessed whether in intact *Arabidopsis* cells proteins follow a melting profile similar to that in lysates when heated (5, 26). To this end, we heated 100-µl aliquots of *Arabidopsis* cells in protein extraction buffer were heated to different temperatures (from 25°C to 80°C) for 2 min and analyzed the lysates by sodium dodecyl-sulfate polyacrylamide gel electrophoresis (SDS-PAGE) after seven freeze-thaw cycles and centrifugation. As expected, the *Arabidopsis* proteins unfolded and precipitated at high temperature (Fig. S2 *B* and *C*).

Subsequently, as a proof of concept, we aimed to apply CETSA to cells treated with bikinin, an inhibitor of several *At*SKs in *Arabidopsis* (12). Application of bikinin at concentrations of 30-50 µM to *Arabidopsis* seedlings induces BR responses that can be measured by changes in the phosphorylation status of the transcription factor BES1/BZR2 (14). To check whether bikinin is effective in *Arabidopsis* cell suspension cultures we treated cells with 50 µM bikinin for 30 min. As expected, bikinin induced the dephosphorylation of BES1 in the cell cultures similarly to the most active BR, brassinolide (BL) (Fig. S2*D*). Taken together, bikinin can induce BR responses in *Arabidopsis* cell suspension cultures.

Next, we investigated the effect of bikinin on the thermal stability of its direct targets, the ten *At*SKs (12), by means of the adjusted CETSA protocol (Fig. S1) and by using Western blots for detection. In brief, after 30 min of treatment with bikinin or dimethylsulfoxide (DMSO), the *Arabidopsis* cells were washed, resuspended in protein extraction buffer containing either bikinin or DMSO, and aliquoted into PCR tubes. Then, the aliquots were heated to 12 distinct temperatures (30, 35, 40, 43, 46, 49, 52, 55,58,61, 65, and 70°C) for 2 min. Afterward, the heated cells were lysed through seven freeze-thaw cycles followed by Western blot-based protein detection. As specific antibodies for all *At*SKs are not available, *Arabidopsis* cell suspension cultures overexpressing the hemagglutinin (HA)-tagged *At*SKs were utilized to generate protein-melting curves and to assess the bikinin-induced T_m_ shifts. First, we examined whether bikinin induced T_m_ shifts for *At*SK12 and *At*SK13 at a 50-µM concentration, but, surprisingly, found that it stabilized *At*SK13 with a T_m_ shift of 5.94°C (Fig. S3*A*), but not *At*SK12 (Fig. S3*B*). By contrast, the thermal stability of the ATP synthaseβ (ATPβ), used as a control, was not affected by bikinin. We further determined the half-maximum effective concentration (EC_50_) of bikinin for *At*SK12 by means of isothermal dose-response fingerprinting (ITDRF_CETSA_) at 45 °C to ensure a sufficient shift in the denaturation temperature. At low concentrations, bikinin did not affect the thermal stability of *At*SK12 and the relative band intensity reached a plateau at approximately 250 µM (Fig. S3*C*). Therefore, to ensure saturation and achieve sufficiently sized T_m_ shifts for all *At*SKs, we used 250 µM bikinin (Fig. 1). Staining of the cells with the cell viability tracer, fluorescein diacetate (FDA), excluded the potential cytotoxic effect of bikinin when used at high concentrations (Fig. S2*E*). Of the 10 putative bikinin targets (12), *At*SK11, *At*SK12, *At*SK13, BIN2/*At*SK21, *At*SK22, and *At*SK41 showed T_m_ shifts, whereas the thermal stability of *At*SK23, *At*SK31, *At*SK32, and *At*SK42 was not affected by the small molecule (Fig. 1) as well as the thermal denaturation of the ATPβ control (Fig. S4). Collectively, these results showed that bikinin stabilized most of its targets, indicating that the CETSA protocol was applicable to intact *Arabidopsis* cells.

**Figure 1.**
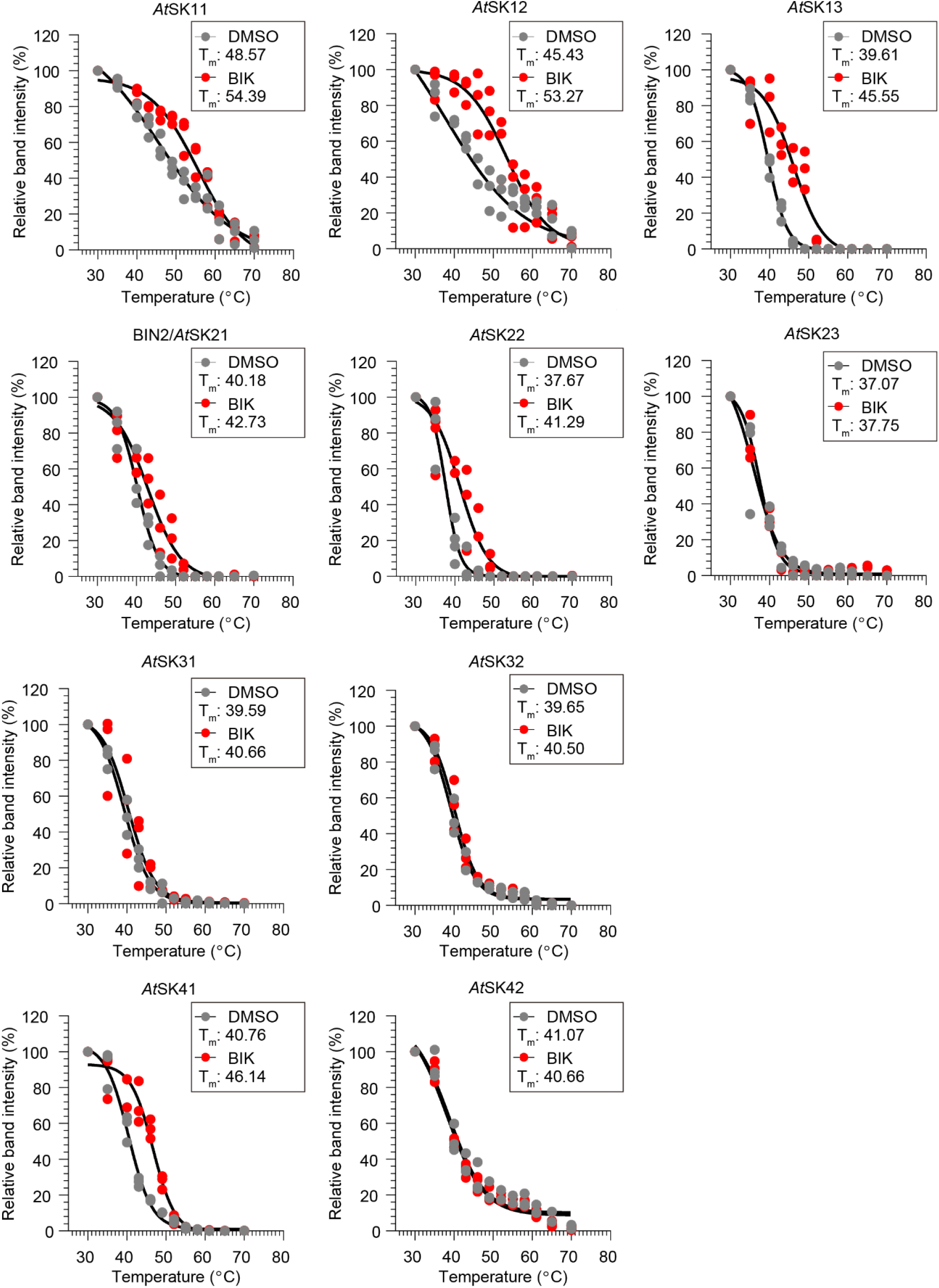
Bikinin stabilized a subset of the *At*SKs. Thermal denaturation curves for 10 hemagglutinin (HA)-tagged *At*SKs stably overexpressed in *Arabidopsis* cell suspension cultures in the presence of 250 μM bikinin (BIK) or 0.1% (v/v) DMSO. The relative band intensities from the Western blot analysis were calculated based on the lowest temperature (30°C). Melting temperatures (T_m_) are indicated. Individual data points are plotted for three biological replicates.

### CETSA MS of bikinin in intact *Arabidopsis* cells

Several recent studies in mammalian cells reported the use of proteome-wide CETSA MS for obtaining a comprehensive view on small molecule-protein interactions through determination of individual temperature shifts (8–10, 27). Therefore, we extended the CETSA protocol as described above (Fig. S1) to the *Arabidopsis* proteome (Fig. S2*D*) by using 50 µM bikinin, the BR response-inducing concentration in cell cultures. Briefly, following the heating at 25, 30, 35, 40, 45, 50, 55, 60, 70, and 80°C and freeze-thaw cycles, samples were analyzed with a nanoscale liquid chromatography coupled to tandem mass spectrometry (nano LC-MS/MS), whereafter they were digested with trypsin and labeled with 10-plex tandem mass tag (TMT10).

In total, 6,000 proteins were identified of which the melting profiles were defined for 4,225 proteins in samples both treated with bikinin and DMSO (Dataset S1*A*). Approximately 96 % of the identified proteins had a melting temperature within the range of 35°C - 60°C (Fig. 2*A* and Dataset S1*A*), of which only 61 proteins displayed a significant change in thermal stability (27 were stabilized and 34 were destabilized) in the presence of 50 µM bikinin (absolute value of T_m_ shift ≥ 2°C, analysis of variance (ANOVA)-based F-test *P* < 0.01) (Fig. 2*B* and Dataset S1). However, because none of the *At*SKs was identified by the CETSA MS, we examined the protein expression of *At*SKs in *Arabidopsis* cell suspension cultures. Although quantitative reverse transcription PCR (qRT-PCR) revealed expression of all *At*SKs in the cell cultures (Fig. S5*A*), only *At*SK11, *At*SK21, *At*SK31 and *At*SK41, albeit at a low intensity were detected with shotgun proteomics (Fig. S5*B* and Dataset S2). Given that all *At*SKs were efficiently extracted using the CETSA protocol when overexpressed (Fig. 1), probably the low number of peptides identifying these proteins in cell cultures obstructed the creation of their melting curves (Dataset S2).

**Figure 2.**
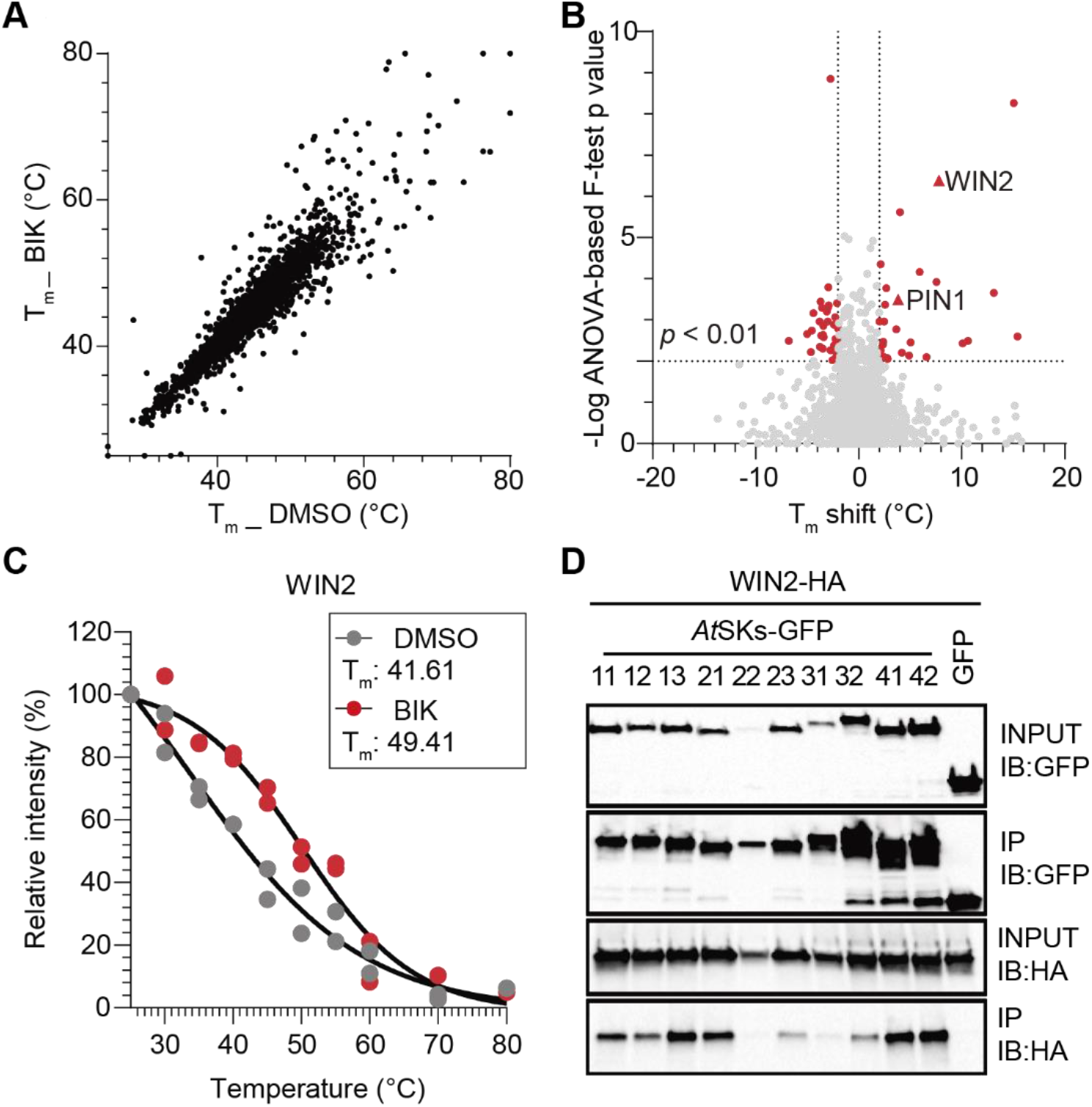
CETSA MS in the presence of bikinin. (**A**) Melting temperatures of the 4,225 proteins identified in the CETSA MS in the presence of 50 µM bikinin (BIK) or 0.1% (v/v) DMSO. (**B**) Distribution of the T_m_ shifts of all proteins. Proteins in red highlight significant changes (T_m_ shift > 2°C, ANOVA-based F-test *P* < 0.01). (**C**) Thermal denaturation curves generated for WIN2 in the presence of 50 μM BIK or 0.1% (v/v) DMSO subtracted from the CETSA MS in *Arabidopsis* cell suspension cultures. The T_m_ is indicated. Individual data points are plotted for *n*=2 technical replicates. (**D**) Coimmunoprecipitation of WIN2 with all *At*SKs when coexpressed in tobacco. The *p35S::WIN2-HA* construct was transiently coexpressed with either *p35S::AtSKs-GFP* or *p35S::GFP* (as a negative control). Proteins were extracted (Input) and immunoprecipitated (IP) by means of GFP beads. *At*SKs-GFP, GFP, and WIN2-HA were detected with anti-GFP and anti-HA antibody, respectively. IP, immunoprecipitation; IB, immunoblot.

As changes in thermal stability had been observed also for downstream effectors of the direct small-molecule target, possibly as a result of altered posttranslational modifications or interactions with other proteins (4, 9), we examined whether any of the 61 proteins (*P* < 0.01) (Dataset S2*B*) functioned together with the *At*SKs and found that only the MITOGEN-ACTIVATED PROTEIN KINASE3 (MPK3); which acts downstream of the known BIN2/*At*SK21 interactor YODA (YDA) (28), exhibited a T_m_ shift (−2.96°C, *P* < 0.01). Furthermore, we checked whether any of the 61 proteins with significant T_m_ shifts were putative *At*SK interactors (Fig. S6) according to the STRING database (29). Based on this analysis, the HOPW1-1-INTERACTING2 (WIN2, AT4G31750) (T_m_ shift = 7.80 °C, *P* < 0.01) (Fig. 2*C*) was selected as a possible *At*SK-interacting protein. Interaction between HA-tagged WIN2 and each of the 10 GFP-tagged *At*SKs was observed by co-immunoprecipitation experiments carried out using tobacco (*Nicotiana tabacum*) cells transiently overexpressing the proteins (Fig. 2*D*). In summary, in cell suspension cultures bikinin affected the thermal stability of the *Arabidopsis* proteome and induced T_m_ shifts in several proteins that might be putative *At*SK-interacting proteins or *At*SK downstream effectors.

### The phosphoproteome of bikinin

As, besides MPK3, the proteins with altered thermal stability identified by the CETSA MS (Dataset S1) were neither direct bikinin targets nor known downstream *At*SK effectors, we carried out a phosphoproteomics analysis on bikinin-treated *Arabidopsis* cell suspension cultures. We hypothesized that the putative *At*SK-interacting proteins or *At*SK downstream effectors might modify their phosphorylation state upon inhibition of the kinase activity of the *At*SKs. *Arabidopsis* cell suspension cultures were treated with 50 µM bikinin or DMSO for 30 min, under the same CETSA MS. The phosphoproteomics analysis revealed that, in total, 9,351 phosphopeptides that were mapped to 1,751 proteins (Dataset S3*A*). Bikinin treatment significantly down regulated (*P* < 0.05) the phosphorylation intensities of 972 phospho-sites that belong to 665 proteins (Fig. 3*A*, Fig. S7 and Dataset S3 *B*-*E*), of which, six proteins were known *At*SK-interacting proteins including, BZR1 (30), ARF2 (21), YDA (28), GLUCOSE-6-PHOSPHATE DEHYDROGENASE6 (G6PD6) (31), TETRATRICOPETIDE-REPEAT THIOREDOXIN-LIKE3 (TTL3) (32), and OCTOPUS (33). In addition, the phosphorylation intensities of 101 phospho-sites belonging to 84 proteins were significantly up regulated (*P* < 0.05) (Fig. 3*A*, Fig. S7 and Dataset S3 *B*-*E*). Of note, both downregulated phosphosites and upregulated phosphosites were identified in 35 proteins (Fig 3*A* and Dataset S3 *B*-*E*). The gene ontology (GO) enrichment analysis (34) showed that most of the enriched terms for the identified phosphorylation-regulated proteins were related to mRNA splicing, metabolic process, and transport (Fig. S8*A*, Dataset S3*G*). However, of the bikinin-regulated phosphoproteins, only six (phosphorylation-downregulated) proteins showed significant T_m_ shifts in the CETSA MS (Dataset S3*E*), including the auxin efflux carrier PIN1 (T_m_ shift = 3.83 °C) (Fig. 3 *A* and *B*). Taken together, only a few proteins identified in the bikinin phosphoproteome displayed bikinin-induced changes in their thermal stability.

**Figure 3.**
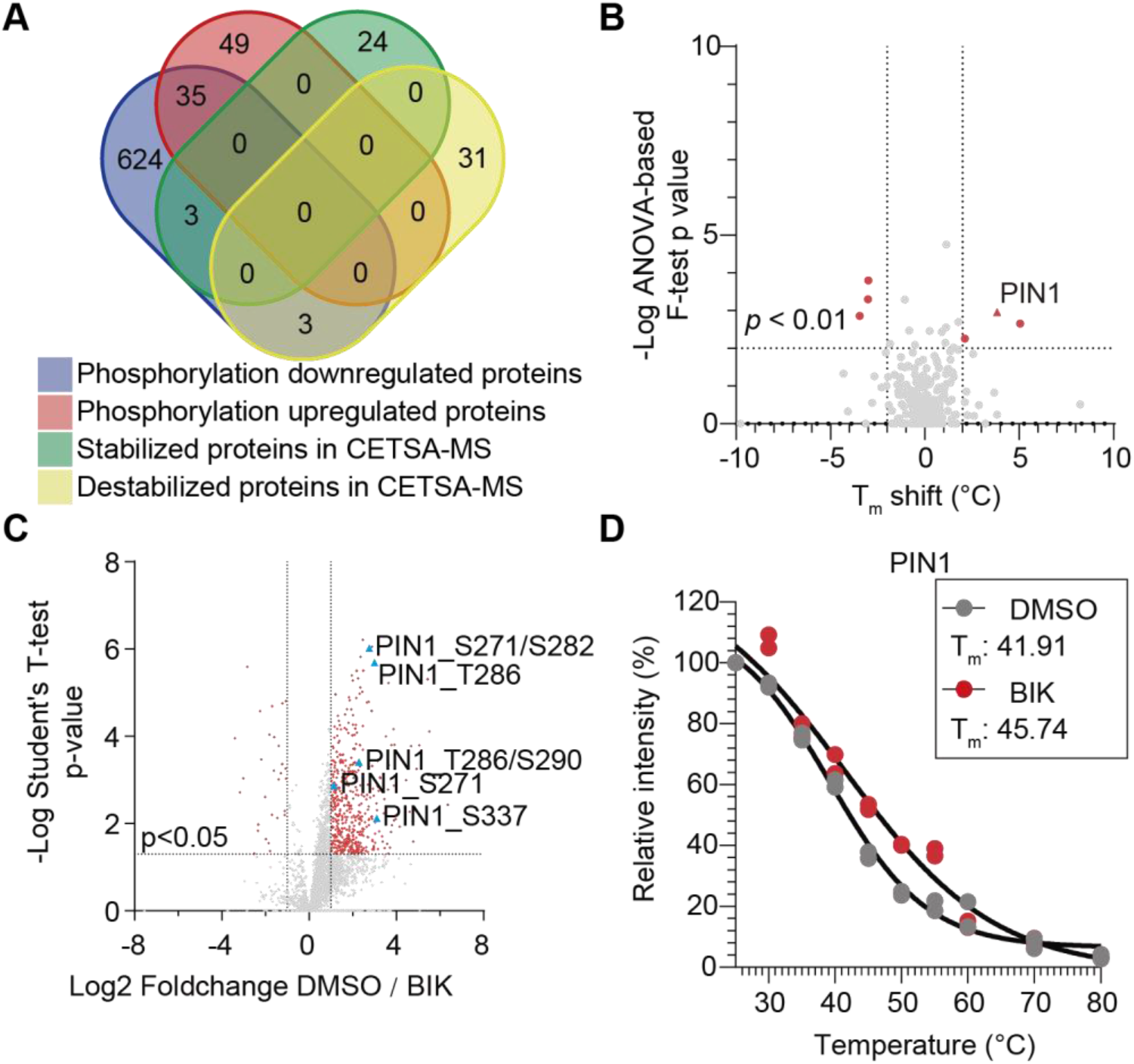
Identification of PIN1. (**A**) Venn diagram comparing the CETSA MS and the bikinin (BIK) phosphoproteome. Only six proteins (phosphorylation downregulated) showed significant T_m_ shifts in CETSA MS. (**B**) T_m_ shifts distribution of the proteins of which the phosphorylation intensities were downregulated in the presence of 50 µM BIK. (**C**) Volcano plot representation of quantitative phosphoproteomics analysis in the presence of 50 µM BIK and 0.1% (v/v) DMSO. Diamonds represent phosphopeptides quantified in five biological replicates, each phosphopeptide log 2 (fold change) is the average logarithmic ratio of phosphopeptide abundance of in cell suspensions treated with BIK vs DMSO plotted against the log 10 (*P* value) determined with the Student’s *t* test. Phosphopeptides of PIN1 are indicated. (**D**) The thermal denaturation curves generated for PIN1 in the presence of 50 μM BIK or 0.1% (v/v) DMSO subtracted from the CETSA MS in *Arabidopsis* cell suspension cultures. The T_m_ is indicated. Individual data points are plotted for *n*=2 technical replicates.

### *At*SKs phosphorylate PIN1 and regulate its polarity

We hypothesized that PIN1 is a direct substrate of the *At*SKs, because the PIN1 protein had been identified in the CETSA MS (Fig. 3 *B* and *D*, and Dataset S1) and the phosphorylation intensities of five residues in the hydrophilic loop (HL) (Ser^271^, Ser^282^, Thr^286^, Ser^290^, and Ser^337^) (Fig. S9) were reduced in the presence of 50 µM bikinin (Fig. 3*C* and Dataset S3*C*). To verify this observation, we carried out a protein kinase assay by incubating the polyhistidine (HIS)-tagged hydrotropic loop (HL) of PIN1 with HIS-small ubiquitin-like modifier (HIS-SUMO)-tagged *At*SK proteins in an *in vitro* phosphorylation reaction. The results showed that eight *At*SK proteins phosphorylated the His-PIN1-HL (Fig. 4*A*). Remarkably, four of the five PIN1 residues, of which the phosphorylation was altered by the bikinin, had previously been identified as targets of the serine/threonine protein kinases PINOID/AGCVIII kinases (PID/WAGs) (35), D6 Protein Kinases (D6PKs) (36), and MPKs (MPK3, MPK4, and MPK6) (37, 38). Moreover, these residues were conserved (Fig. S9) and essential for the polar localization of the PIN proteins with long HLs (39).

**Figure 4.**
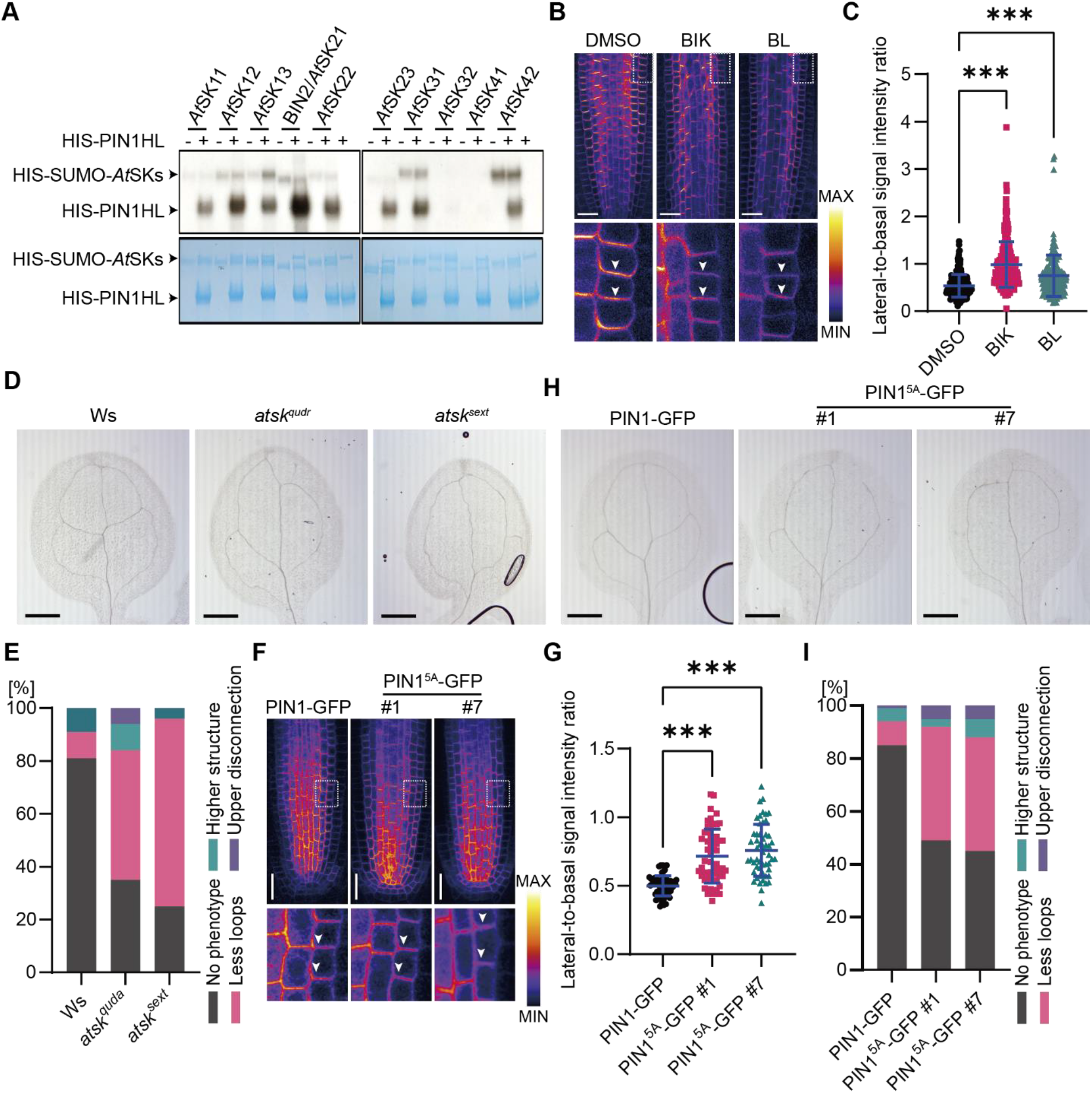
PIN1 phosphorylation and polarity modulation by the *At*SKs. (**A**) Phosphorylation of HIS-PIN1HL by HIS-SUMO-*At*SKs *in vitro*. The signal of the *At*SKs-induced phosphorylation of HIS-PIN1HL were indicated in the autoradiography (Top). The Coomassie Brilliant Blue (CBB) showed the presence of the respective recombinant proteins (bottom). (**B**) Immunolocalization of PIN1 in root tips after 12 h of 50 µM bikinin (BIK), 10 nM brassinolide (BL), or 0.1% (v/v) DMSO treatments. The pictures below each main image represent zoomed in regions. Scale bars, 20 µm. (**C**) Quantitative evaluation of (**B**) showing mean of PIN1 lateral-to-basal signal intensity ratio in endodermal cells. *n* > 180 cells corresponding to a minimum of 15 roots per treatment from three independent experiments. (**D**) Representative images of venation patterning defects in cotyledons of Wassilewskija (Ws), *At*SK quadruple mutant (*At*SKs^qudr^), and sextuple mutant (*At*SK^sext^). Scale bars, 1 mm. (**E**) Quantification of venation defects in *At*SK^qudr^ and *At*SK^sext^ mutants (*n* > 60 of each genotype from three independent experiments). (**F**) Subcellular localization of PIN1-GFP and PIN1-GFP^5A^ in root tip cells. The pictures below each main image represent zoomed regions. Arrowheads indicate the lateral membrane signal in the endodermal cells. (**G**) Quantitative evaluation of (**C**) showing mean of PIN1-GFP and PIN1-GFP^5A^ lateral-to-basal signal intensity ratio in endodermal cells. *n* > 60 cells corresponding to a minimum of 15 roots per genotype from three independent experiments. (**H**) Representative images of venation patterning defects in cotyledons of *PIN1pro:: PIN1-GFP* and two transgenic lines of *PIN1pro::PIN1*^*5A*^*-GFP*. Scale bars, 1 mm. (**I**) Quantification of venation defects in *PIN1pro:: PIN1-GFP* and two *PIN1pro::PIN1*^*5A*^*-GFP* transgenic lines (*n* > 60 of each genotype from three independent experiments). (**C** and **G**) Scatter dot plots showing all the individual points with the means and standard errors. One-way ANOVA with Tukey’s post hoc test compared to DMSO or PIN2-GFP respectively, \*\*\**P* < 0001.

Given the prominent role of phosphorylation in regulating polarity, we next assessed whether treatments with bikinin and BL affected the polar localization of PIN1 and PIN2 in the root meristem by means of immunolocalization. Wild type *Arabidopsis* plants were treated in liquid medium with 50 µM bikinin, 10 nM BL, and 0.1% (v/v) DMSO for 12 h. As anticipated, PIN1 displayed a more apolar localization in the presence of bikinin and BL (Fig. 4 *B* and *C*) than with DMSO, whereas the PIN2 polarity remained unaffected (Fig. S10). Given that PIN1-mediated polar auxin transport regulates the foliar vascular patterning (40), we examined whether *At*SKs were involved in the regulation of leaf venation. Five-day-old wild type *Arabidopsis* plants, germinated and grown in liquid medium containing 50 µM bikinin and 10 nM BL, had an abnormal vascular patterning in the cotyledons, mainly with missing loops, when compared to the mock control (Fig. S11). To provide genetic evidence for *At*SK function in leaf venation, we analyzed the vascular pattern of the *At*SK quadruple (*atsk13RNAi bin2 bil1 bil2*, designated *atsk*^*quad*^) and sextuple (*atsk11RNAi atsk12RNAi atsk13RNAi bin2 bil1 bil2*, designated *atsk*^*sext*^) mutants (41). Consistent with BR and bikinin treatments, the *atsk*^*quad*^ mutant exhibited an abnormal vascular patterning, with missing loops and disconnected upper loops (Fig. 4 *D* and *E*). Because of the seedling lethality of the *atsk*^*sext*^ mutant (41), the plant phenotypes were analyzed in the T1 generation. The *atsk*^*sext*^ mutant also displayed an abnormal vascular patterning, but mainly with missing loops (Fig. 4 *D* and *E*).

To test whether the identified five phosphorylation sites were relevant for the *At*SK-mediated regulation of PIN1, we generated phospho-inactive His-PIN1-HL^5A^ by substituting the Thr and Ser residues with Ala. The phosphorylation of His-PIN1-HL^5A^ by BIN2/*At*SK *in vitro* was reduced but not abolished (Fig. S12*A*), suggesting that more sites were phosphorylated by BIN2/*At*SK21. Next, we introduced the phospho-inactive PIN1^5A^-GFP (*PIN1pro::PIN1*^*5A*^*-GFP*) into wild type *Arabidopsis* plants. Noteworthy, eight of 21 GFP-expressing transgenic plants had naked inflorescence stems in the first-generation (Fig. S12*B*), reminiscent of the *pin1* null mutant (42). Then, we examined the localization of PIN1^5A^-GFP in the root tip cells of wild type-resembling transgenic plants. When expressed at the same levels as the native PIN1-GFP (*pPIN1::PIN1-GFP*) (43) (Fig. S12*C*), PIN1^5A^-GFP exhibited a more apolar localization in two independent transgenic lines (Fig. 4 *F* and *G*) that, similarly way to the *atsk*^*quad*^ and *atsk*^*sext*^ mutants, had an abnormal vascular patterning in the cotyledons (Fig. 4 *H* and *I*). In summary, these observations show that *At*SK-mediated phosphorylation of PIN1 is important for their polar localization and for leaf venation.

## Discussion

Given the challenges in small-molecule target identification in plants (44), novel experimental strategies, especially label-free methods, are required to facilitate chemical genetics studies. CETSA is one such method that has been proven useful for the detection of direct drug targets and downstream effects of drug-induced perturbations in several cellular systems and tissues (9, 27, 45, 46). In plants, CETSA has been successfully applied for small-molecule target validation in cell lysates of *Arabidopsis* (6) and, recently, the thermal profiles of more than 2,000 proteins in *Arabidopsis* lysates have been reported (5, 26). Here, we explored the potential of CETSA to identify the protein targets of the *At*SK kinase inhibitor bikinin (12) in intact *Arabidopsis* cells. In Western blot-CETSA, bikinin stabilized six of the ten *At*SKs. As expected, bikinin had no effect on either *At*SK31 or *At*SK42, of which the kinase activity was not inhibited *in vitro* (12), but surprisingly, on *At*SK23 and *At*SK32 as well, of which the *in vitro* inhibition of the kinase activity had previously been reported (12). One reason might be that the binding affinity of bikinin to *At*SK23 and *At*SK32 is lower than that to other *At*SKs, thus requiring higher bikinin concentrations to induce the noticeable T_m_ shifts. Moreover, in a previous study, approximately 30% of the target proteins of the promiscuous kinase inhibitor staurosporine did not show T_m_ shifts (9), suggesting that some protein kinases are not responsive in CETSA.

By combining CETSA with MS, we identified the thermal profiles of 4,225 proteins in the presence of bikinin in *Arabidopsis* cell suspension cultures, of which 61 (1.44%) showed significant T_m_ shifts (absolute value ≥ 2°C, ANOVA-based F-test *P* < 0.01). Although the CETSA MS assay did not detect any *At*SKs, probably due to their low protein abundance in the cell cultures, MPK3, the substrate of the know BIN2/*At*SK21 interactor YDA (28), was identified. In addition, the thus far unknown interaction between WIN2 (AT4G31750) and *At*SKs was validated *in vivo*. WIN2 is implicated in the modulation of the defense responses of induced by the *Pseudomonas syringae* effector protein HopW1-1 (47). *At*SK11 has been shown to regulate the pattern-triggered immunity and the susceptibility to *P. syringae*, probably through G6PD6 phosphorylation (48). Our data suggest that *At*SK11, as well as its homologs, might regulate the immune responses to *P. syringae via* direct interaction with WIN2.

To demonstrate the capability of CESTA MS to identify proteins that might bind the *At*SKs directly or function in downstream pathways, we carried out a phosphoproteomics analysis in the presence of bikinin. Phosphorylation sites in 665 proteins were downregulated upon bikinin treatment, comprising six known substrates of the *At*SKs, hence, confirming the quality of our data. Noteworthy, the phosphorylation of 84 proteins was upregulated upon bikinin treatment, indicating that these proteins might be indirect *At*SK targets. The GO enrichment analysis for the bikinin-regulated phosphoproteins revealed that *At*SKs might be involved in the regulation of RNA splicing and protein intracellular transport, like their mammalian homologs (49, 50). Moreover, of all proteins with differential phosphorylation, 76 proteins were also identified by proximity labelling with BIN2/*At*SK21 as a bait (51) (Dataset S3*F*). Thus, we speculate that some of these proteins might be novel substrates of the *At*SKs. By comparing the CETSA MS and phosphoproteome datasets, we found that the thermal stability of most of these proteins was not affected, even though their phosphorylation intensities were altered by the bikinin treatment. A possible reason might be that either the phosphorylation cannot affect the protein melting behavior or that the in the bikinin-induced changes in the phosphorylation intensity are not sufficient to induce an important T_m_ shift for most of these proteins. However, for a few proteins, bikinin had an impact on both the phosphorylation and melting behaviors, indicating that the phosphorylation status might affect the thermal stability of some proteins. Although some studies in human cells reported that phosphorylation affected the protein thermal stability (52), others demonstrated that for most of the proteins, the melting behavior of phosphorylated and non-phosphorylated forms were concordant (53, 54).

Particularly, bikinin reduced the phosphorylation and induced the thermal stabilization of the auxin efflux carrier PIN1. The identified phosphorylation sites in the HL of PIN1 by *At*SKs are conserved among the PIN proteins (Fig. S9), some of which are essential for their polar localization and the intercellular auxin transport (39). However, although some kinases shared phosphorylation sites in the PIN proteins, for instance PID/WAGs and D6PKs, their corresponding mutants or transgenic overexpression lines affected the PIN polarities differently (35, 36). Moreover, the auxin-regulated receptor CANALIZATION-RELATED AUXIN-REGULATED MALECTIN-TYPE RECEPTOR-LIKE KINASE (CAMEL) (55) and CALCIUM-DEPENDENT PROTEIN KINASE 29 (CPK29) (56) phosphorylate PIN1 and controls its polarity via phosphorylation sites that are not shared with other kinases, including PID/WAGs, D6PKs, MPKs and the *At*SKs (Fig. S9) (55, 56). Therefore, multiple parallel mechanisms for the maintenance of the PIN polarities and activities exists. Our data imply that BRs control the PIN1 phosphorylation and polarity *via At*SKs. Two of the five phosphorylation sites identified in our phosphoproteomics study are also targeted by PID/WAGs and D6PKs, whereas another two phosphorylation sites are targeted by MPK3, MPK4, and MPK6, implying a functional redundancy of *At*SKs with other kinases in the PIN phosphorylation. In addition, bikinin and BR treatments, which inhibit the activity of *At*SKs directly and indirectly, respectively, impaired the PIN1 polarity, indicating that *AtS*Ks control the polarity of PINs in combination with other kinases. Considering this overlap in the PIN1 phosphorylation sites, understanding of the precise environmental or developmental conditions and upstream pathways that coordinate these kinases will be needed to cooperatively control the PIN polarities. Furthermore, because the PIN phosphorylation induced by D6PKs and PID/WAGs control their activity (57), it will be necessary to examine whether bikinin and exogenous BRs disrupt the auxin transport activity of PIN1 and other PINs. Previous studies have shown that BRs affect PIN proteins through different mechanisms (23, 58), including transcriptional activation of of *PIN4* and *PIN7* (58). Additionally, BRs and bikinin stabilized PIN2, but not PIN1, to respectively interfere with the auxin distribution in gravistimulated roots (23) and BRs partially altered PIN2 polarity via the actin cytoskeleton (59). However, others demonstrated that the actin cytoskeleton was not required for the polar localization of PIN2 (60). Interestingly, BRs did not affect the PIN2 polarity in our study, although four of the five identified phosphorylation sites are conserved in the HL of PIN2. Together, these data indicate that the mechanisms controlling the polarity and the stability of PINs by BRs differ.

PIN1 has been shown to be expressed in early leaf veins and margins and PIN1-mediated polar auxin transport to be important for the leaf venation pattern in *Arabidopsis* (40). Moreover, MPK6 could regulate the leaf venation pattern probably by regulating the PIN1 phosphorylation and polarity, although the main MPK6-targeted phosphorylation site Ser^337^, also targeted by *At*SKs, was not involved (37). In addition, the auxin-regulated receptor CAMEL controls the cotyledon venation through modulation of the PIN1 phosphorylation and respectively polarization (55). Our data showed that chemical activity inhibition or genetic knockout of the *At*SKs resulted in defective cotyledon venation pattern, suggesting that BRs control leaf venation via *At*SK-mediated phosphorylation of PIN1. It remains to be established whether the PIN1 polarity in the veins is affected under these conditions.

In summary, we adapted the CETSA MS method for intact *Arabidopsis* cells and demonstrated its ability to identify direct targets and downstream components of the small-molecule target proteins. Such information is useful to understand the mode of action of the small molecules and the function of their target proteins. Moreover, we identified PIN1 as a novel substrate of *At*SKs, uncovering an unknown BR mechanism and auxin crosstalk.

## Materials and Methods

Details on the material and experimental procedures used in this study (i.e., plant material, CETSA and CETS MS assays, phosphoproteomics, generation and transformation of constructs, confocal microscopy, *in vitro* kinase assay, whole-mount *in situ* immunolocalization of PIN1, immunoprecipitation and statistical analysis) can be found in *SI Materials and Methods*.

## Acknowledgments

We thank Yanhai Yin for providing the anti-BES1 antibody, Johan Winne and Brenda Callebaut for synthesizing bikinin, Yuki Kondo and Hiroo Fukuda for published materials, and Martine De Cock for help in preparing the manuscript.

