## Supplementary Information for "Proteome-wide cellular thermal shift assay reveals novel crosstalk between brassinosteroid and auxin signaling"

**This PDF file includes:**

Supplementary text
Figures S1 to S12
Tables S1
SI References

**Other supplementary materials for this manuscript include the following:**

Datasets S1 to S3

### Supplementary materials and methods

#### Plant materials and growth condition

*Arabidopsis thaliana* (L.) Heynh. lines with in the Columbia-0 (Col-0) accession were used, except the *atsk13RNAibin2-3bil1bil2* quadruple and *atsk11RNAiatsk12RNAiatsk13bil2-3bil1bil2* sextuple mutants were generated with the Wassilewskija (Ws-0) accession as previously described (1). The transgenic *Arabidopsis* plant expressing *PIN1pro::PIN1-GFP* had been described previously (2).

The *Arabidopsis* seeds were stratified for 2 days at 4°C, germinated, and grown on half-strength Murashige and Skoog (½MS) agar (1% w/v) plates containing 1% (w/v) sucrose at 22°C with a 16-h light/8-h dark cycle for 5 days under 120 µmol $m^{-2}$ $s^{-1}$ of photosynthetically active radiation. *Arabidopsis* PSB-L cell suspension culture was cultured in MS basal salts with minimal organics (MSMO) medium at 22°C, a 16-h light/8-h dark photoperiod under 60 µmol $m^{-2}$ $s^{-1}$ of photosynthetically active radiation and gentle agitation (130 rpm).

#### Constructs generation

The genomic fragment of the *WIN2* gene without stop codons was amplified with the attB sites compatible with the Gateway system. The PCR products were then introduced into the *pDONR221* donor vectors (Invitrogen). The entry clones *pDONRP4-P1R-p35S*, *pDONR221-AtSKs* (3), *pDONR221-WIN2*, and *pDONRP2R-P3-HA* were recombined in a multisite LR reaction with *pH7m34GW* (Invitrogen) as the destination vector. For the constructs used for protein purification, the coding sequence (CDS) fragments of the 10 *At*SKs were amplified and introduced into a *pET-SUMO* vector (4) by means of the Gibson cloning method. *pET28a-PIN1HL* (5) and *pDONR221-PIN1-GFP* (6) were used as templates to generate the *pET28a-PIN1HL$^{5A}$* and *pDONR221-PIN1$^{5A}$-GFP* constructs by using one-step site mutation PCR (7), respectively. The entry clones *pDONRP4-P1R-pPIN1* (6) and *pDONR221-PIN1$^{5A}$-GFP* were recombined in a multisite LR reaction with *pB7m24GW,3* (Invitrogen) as the destination vector. The resulting construct *pPIN1::PIN1$^{5A}$-GFP* was introduced into Col-0. Primers used to generate the constructs are listed in Table S1. All clones were confirmed by sequencing.

#### Cell suspension culture transformation

The generated constructs were used to transform dark-grown *Arabidopsis* PSB-D cell suspension cultures as described previously (8). In brief, the *Agrobacterium tumefaciens* culture was washed three times and resuspended in MSMO medium until an $OD_{600}$ = 1.0. Then, 3 ml of the cell suspension culture and 200 µl of washed agrobacteria were cocultured and incubated with 200 µM acetosyringone for 2 days in the dark at room temperature with gentle agitation (130 rpm). Two days after cocultivation, the transformed cell suspension culture was selected by adding 7 ml of MSMO medium containing 500 µg/ml carbenicillin, 500 µg/ml vancomycin and, 20 µg/ml hygromycin. The stable transgenic cell suspension cultures were selected in antibiotics mixture-containing MSMO medium 11 and 18 days after cocultivation.

**Chemical treatments**

Stock solutions of bikinin (50 mM) (homemade) and brassinolide (20 µM) (FUJIFILMWako Chemicals) were prepared in DMSO. The aliquots were stored at -20°C. All chemical treatments were done at room temperature in growth medium and compared with control samples incubated with equal volumes of solvent.

**Microscopy**

For vasculature analysis the cotyledon, 5-day-old seedlings were kept in ethanol overnight to remove chlorophyll followed by a 30-min incubation in 90% (v/v) ethanol and 10% (v/v) acetic acid at 60° and a 2-h incubation in 50% (v/v) ethanol and 625 mM NaOH at 60°C. Finally, the seedlings were incubated in chloral hydrate-saturated lactic acid before monitoring with a bright-field binocular microscope (Leica) with differential interference contrast (DIC) optics and a DXM1200C camera.

The localization of PIN1-GFP and PIN1$^{5A}$-GFP was analyzed with a SP8 confocal microscope (Leica) in root tips of 5-day-old transgenic plants. Images were captured at 488 nm laser excitation and 495–530 nm emission. PIN1 polarity was analyzed quantitatively as described (9)

**Cellular thermal shift assay-Western blot**

Four days after subcultivation, 30 ml of *Arabidopsis* cell suspension culture expressing HA-tagged *At*SKs were treated with 0.1% (v/v) DMSO or 50 µM or 250 µM (as indicated) bikinin for 30 min in MSMO medium. Then, the cell suspension cultures were centrifuged at 300×*g* for 3 min and the supernatants were removed avoiding cell perturbation. Next, the cells were washed with buffer containing 25 mM HEPES (Sigma-Aldrich), 25 mM NaCl, and 0.1% (v/v) DMSO or  bikinin (50 µM or 250 µM). Afterward, 2 ml of buffer containing 25 mM HEPES (Sigma-Aldrich), 25 mM NaCl, cOmplete™ Protease Inhibitor Cocktail tablet (Roche), and 0.1% (v/v)DMSO or bikinin (50 µM or 250 µM) were added, whereafter the supernatants were removed. Subsequently, the cells were distributed into PCR tubes. To ensure the presence of enough proteins, two PCR tubes were prepared with 100 µl cells each for each temperature. Then, the PCR tubes were heated at 12 different temperatures (30, 35, 40, 43, 46, 49, 52, 55, 58, 61, 65, and 70°C) for 2 min in an Applied Biosystems Veriti Thermal Cycler (Thermo Fisher Scientific). After cooling down at room temperature for 2 min, the samples were frozen in liquid nitrogen. To extract the proteins, the samples were thawed at 20°C for 1 min and frozen in liquid nitrogen again. After 7 cycles of freezing and thawing, the samples were homogenized with two metal balls and a Restch mixer mill and centrifuged at 15,000×*g* twice to collect the supernatants. The supernatants were mixed with the required volume of 4× NuPAGE LDS sample buffer (Invitrogen) and 10× NuPAGE sample reducing agent (Invitrogen), heated at 70°C for 10 min, and loaded onto 4-20% Mini-PROTEAN TGX precast gels. The proteins were transferred to polyvinylidene difluoride (PVDF) membranes by means of the Trans-Blot® Turbo™ Transfer System (Bio-Rad). The membranes were probed with the antibodies anti-HA- horseradish peroxidase (HRP) (1:10000; Abcam ab1190) and rabbit-ATPβ (1:2000; Agrisera). The secondary antibodies were ECL™ anti-rabbit IgG, (HRP)-linked whole antibody (GE Healthcare). The blots were developed with Western Lightning Plus-ECL, Enhanced Chemiluminescence Substrate (Perkin-Elmer), and imaged with a ChemiDoc XRS+molecular imager (Bio-Rad). Intensities of protein bands were measured with the Bio-Rad Image Lab software package. The ratio of different temperatures to the lowest temperature (30°C) for bikinin- and DMSO-treated samples was calculated.

**Cellular thermal shift assay – mass spectrometry**

***Sample preparation***: Similar with the Western blot-based CETSA, after 50 µM bikinin or 0.1% (v/v) DMSO treatments and washing, the wild-type *Arabidopsis* cells were heated at 10 different temperatures (25, 30, 35, 40, 45, 50, 55, 60, 70, and 80°C). Following the freeze-thaw cycles and collection of supernatants, the samples were digested with trypsin to generate the peptides. After approved digest, half the volume (corresponding to 50 µg) of each peptide sample was labeled for 3 h at room temperature with 10-plex Tandem Mass Tag reagents (TMT10, Pierce). The treated and control samples were labeled in parallel with the same specific TMT label for the same temperature fraction in each set. When labeled, the samples were quenched by addition of 50 µL 1 M Tris, pH 7.4. The compound and vehicle samples were combined separately in 1 mL 0.5% (v/v) formic acid and subsequently acidified with addition of 200 µL of 10% (v/v) formic acid and desalted with a spin cation exchanger (Strata-XC, Phenomenex). The samples were eluted in 5% (v/v) ammonia in 30% (vv) methanol. Liquid chromatography–mass spectrometry (LC–MS) grade liquids and low-binding tubes were used throughout the purification. The samples were dried in a centrifugal evaporator and, subsequently, further purified with the SP3 methodology (10), before being dissolved in 20 mM ammonia in high-performance liquid chromatography (HPLC) grade water and subjected to HPLC prefractionation thorugh a high pH reversed-phase approach (A buffer, 20 mM ammonia; B buffer, 20 mM ammonia in 80% [v/v] acetonitrile)with a pH-stable reverse-phase column (Zorbax 300 Extend C-18, 4.6 mm × 250 mm; Agilent Technologies) and liquid chromatography AKTA Micro (GE-Healthcare) system. The prefractionation fractions were combined in a concatenated fashion to yield 10 final pooled fractions per sample.

***LC-MS/MS analysis.*** The 10 final fractions from the prefractionation (basic high pH reversed-phase HPLC fractionation) were evaporated to dryness and resuspended in 15 µL of HPLC buffer A (3% [v/v] acetonitrile, 0.1% [v/v] formic acid). An Ultimate 3000 RLSCnano system interfaced to a Q Exactive HF OrbiTrap instrument (Thermo Fisher Scientific) was used for the nanoLC-MS/MS data collection. The nanoLC-MS interface operated with the EasySpray source format fitted with a 50-cm analytical column (<2 µm particles, P/N ES803; Thermo Fisher Scientific) and a 75 µm × 20 mm trap column with 3 µm 100-Å C18 particles (Acclaim PepMap 100; Thermo Fisher Scientific). The elution buffer (B) was 90% [v/v] acetonitrile in water with 0.1% [v/v] formic acid and 5% [v/v] DMSO (all solvents and solvent additives were of HPLC grade or better). Peptides were eluted over a 115-min gradient (3 – 42% buffer B). NanoLC-MS/MS data were obtained with a higher-energy collisional dissociation (HCD) fragmentation on the QE HF instrument by means of a Top5 strategy. Full MS data were assembled with a 60 K resolution up to a maximal signal of 3e6. MS/MS spectra (60 K resolution) were collected at a normalized collision energy of 33 with an isolation width of 1.2 *m/z*, for a maximal injection time of 200 ms or to a maximal ion signal of 1e5 to avoid coalescence. All prefractionated samples were injected and analyzed twice on the LC-MS/MS with 40% of the material injected on-column for each replicate.

***Protein identification and TMT-based quantification.*** Proteins were identified by a database search against 31,360 *Arabidopsis thaliana* protein sequences in Uniprot (Uniprot reference Proteome ID: UP000006548, download date: 2016-10-18) with the Sequest HT algorithm as implemented in the ProteomeDiscoverer 2.1 software package. Search tolerance setting included a mass accuracy of 25 ppm and 50 mDa for precursor and fragment ions, respectively. Noise was reduced by applying a Top N peaks filter, allowing only the 12 strongest fragment ions per 100 Da. A maximum of two cleavage sites were acceptable with the full tryptic cleavage enzyme specificity (K\, R\, no P). The allowed dynamic modifications were carbamidomethylation of Cys, oxidation of Met, and deamidation of Asn and Gln as well as those of protein N-termini by acetylation. TMT modifications of Lys and of peptide N-termini were set as static. The protein identification was validated at the peptide-spectrum-match (PSM) level with the following acceptance criteria; 1% false discovery rate (FDR) determined by Percolator scoring based on Q-value, rank 1 peptides only, and Xcorr ≥ 2.0. For quantification, a maximum co-isolation of 50 % was allowed and a minimum average reporter S/N threshold of 10 was set. Peptides used for quantification included unique and so-called "Razor" peptides. Reporter ions were integrated at a 10-ppm tolerance and verified by manual inspection to ensure that the tolerance setting was applicable.

***Statistical analysis.*** To estimate effect size and *P* value (significance) of the protein stability changes in the melt curve (MC) CETSA MS experiments, the individual protein melting curves were fitted via a nonlinear least squares algorithm with the formula:

$$FC(T)=(1-pl)/(1+ e^{((1-T_m/T)/b)})+pl$$

where the fold change (FC), the measured protein fold change is at temperature T relative to the lowest temperature point; pl, high-temperature plateau of the melting curve; $T_m$, melting temperature; b, parameter corresponding the melting curve slope. The significance of the compound-induced protein thermal stability change was assessed by ANOVA-based F-test comparing the curve fits to the trivial model.

**Co-immunoprecipitation experiments**

*Agrobacterium* strain C58, carrying the constructs of *35S::AtSKs-HA*, *35S::WIN2-GFP*, or *35S::GFP* (as negative control) were coinfiltrated with a p19-harboring strain in the abaxial side of *Nicotiana benthamiana* (tobacco) leaves. After 48 h of infiltration, proteins were isolated with extraction buffer (20 mM Tris-HCl, pH 7.5, 150 mM NaCl, 10 mM DTT, 1% [v:v] NP-40, 1 cOomplete protease inhibitor; [Sigma-Aldrich]) in a 1:2 (w/v) ratio. The lysates were incubated with GFP-Trap magnetic agarose beads (Chromotek) for 2 h at 4°C. The beads were collected with a DynaMag™-2 Magnetic separation rack and washed three times with washing buffer (20 mM Tris-HCl, pH 7.5, 150 mM NaCl, 0.5% [v/v] NP-40). The enriched proteins were released from the beads by boiling in NuPAGE™ LDS Sample Buffer and analyzed by Western blot with α-GFP-HRP and α-HA-HRP antibody.

**Phosphoproteomics**

***Sample preparation.*** Phosphoproteomics were conducted as previously described (11). In brief, 4 days after subcultivation, 30 ml *Arabidopsis* cell suspension cultures were treated with 0.1% (v/v) DMSO or 50 µM bikinin for 30 min in MSMO medium and five independent biological replicates were collected for each treatment. Then, for each of them, 500 mg of ground cells were collected and homogenized in 5 ml protein extraction buffer (50 mM Tris-HCl, pH 8, 0.1 M KCl, 5 mM EDTA, 500 mM DTT, 30% [v/w] sucrose and Milliq water in 50 ml, with 1 tablet Complete Ultra EDTA-free Protease Inhibitor Cocktail Tablet [Roche], 1 tablet Phosphatase Inhibitor Cocktail Tablet PhosSTOP [Roche]). The samples were sonicated on ice and centrifuged at 2500×*g* at 4°C for 15 min to remove debris. Supernatants were collected and 15 ml methanol, 5 ml chloroform, and 20 ml water were added and shaken vigorously; then samples were centrifuged at 5000×*g* for 10 min and the aqueous phase was removed. Subsequently, 20 mL methanol added to the bottom phase remaining in each tube and mixed. The proteins were pelleted via centrifugation at 2500×*g* for 10 min. Pellets were washed with 80% [v/v] acetone and resuspended in 6 M guanidinium hydrochloride in 50 mM triethylammonium bicarbonate (TEAB) buffer (pH 8). Cysteines were alkylated by addition of a combination of tris(carboxyethyl)phosphine (TCEP; Pierce) and iodoacetamide (Sigma-Aldrich) to final concentrations of 15 mM and 30 mM, respectively. The reaction was allowed for 15 min at 30°C in the dark, whereafter 8 M urea was added. Of each sample 3 mg was predigested with EndoLysC (Wako) in 1:100 (w:w, 1 aliquot of 10 µg) at 37°C, and mixed for 2.5 h in the dark. Then the samples were diluted 8× with 50 mM TEAB, followed by a trypsin digestion overnight (Trypsin Gold, mass spectrometry grade; Promega) at 37°C and at an enzyme-to-substrate ratio of 1:100 (w:w). Finally, the solution was adjusted to pH ≤ 3 with trifluoroacetic acid (TFA) to arrest digestion and desalted with SampliQ C18 SPE cartridges (Agilent) according to the manufacturer's guidelines. The dried eluates were resuspended in 500 µl of loading buffer (80% [v/v] acetonitrile, 6% [v/v] TFA) and incubated with 1 mg of MagReSyn Ti-IMAC microspheres (ReSyn Biosciences) for 20 min at room temperature. The supernatant was removed and 500 µl of loading buffer was added for 30 s to remove unbound samples. The microspheres were washed once with 500 µl wash buffer A (60% [v/v] acetonitrile, 1% [v/v] TFA, 200 mM NaCl) for 2 min and then twice with 500 µl wash buffer B (60% [v/v] acetonitrile, 1% [v/v] TFA). The bound phosphopeptides were eluted with adding 80 µl elution buffer (40% [v/v] acetonitrile, 1% [v/v] $NH_4OH$) for 15 min and repeated 3 times. Then, 6 µl of 100% [v/v] formic acid was added to the first 80-µl eluate to acidify the solution. This elution was repeated twice for a final elution volume of 240 µl. The microspheres were cleared and the phosphopeptide-containing eluate was transferred to a new tube. Samples were vacuum-dried and then subjected to LC-MS/MS analysis (11).

***Database search and data analysis.*** MS/MS spectra were searched against the Arabidopsis Information Resource (TAIR) 10 database for *Arabidopsis thaliana* (version TAIR10 _pep_20101214) by the MaxQuant software (version 1.5.4.1). MaxQuant settings can be found in the Supplementary Information (Dataset S3*E*). The 'Phospho(STY).txt' output file generated by the MaxQuant search was loaded into the Perseus software (version 1.6.10.45) for analysis. All the data were selected firstly by a localization prob-cut-off >0.75. Log2-transformed phosphosite label-free quantification (LFQ) intensities were used for further analysis and the two-sample test with a *P* value cut-off <0.05 was carried out to test for differences among the treatments.

#### *In vitro* kinase assay

Recombinant HIS-SUMO-*At*SKs and His-PIN1HL were incubated in kinase reaction buffer (50 mM Tris-HCl, pH 7.5, 100 mM NaCl, 10 mM $MgCl_2$, and 10 µM adenosine 5'-triphosphate) in the presence of 5 µCi [$\gamma$-$^{32}$P]-ATP (NEG502A001MC; Perkin-Elmer) at 25°C for 60 min. The reactions were terminated by adding NuPAGE LDS sample buffer (Invitrogen) and NuPAGE sample reducing agent (Invitrogen), separated by 4-20% sodium dodecyl sulfate-polyacrylamide gel electrophoresis (SDS-PAGE) and stained with Coommasie Brilliant Blue. Gels were dried and radioactivity was detected by autoradiography on a photographic film with a FLA 5100 phosphor imager (Fujifilm).

#### Real-Time Quantitative Reverse Transcription PCR

Total RNA was extracted from 100 mg plant material with with TRIzol (Invitrogen), followed by on-column purification with the RNeasy mini kit (Qiagen). cDNA was generated with the iScript cDNA synthesis kit

(Bio Rad). *ATSK* and *EF1a* genes were amplified from 1,000 ng total RNA using SYBR green I qPCR master mix (Roche) and LightCycler 480 (Roche). All primers are listed in Table S1.

**Whole-mount *in situ* immunolocalization of PIN1**

PINs were immunolocalized in primary roots as described (12). In brief, 4-day-old Col-0 seedlings were treated with 50 µM bikinin, 10 nM BL, or 0.1% (v/v) DMSO in liquid ½MS medium for 12 h. The anti-PIN1 and anti-PIN2 antibodies were used at a 1:1000 dilution. The secondary goat anti-rabbit antibody coupled to Cy3 (Sigma-Aldrich) was diluted 1:600. Samples were imaged with a LSM800 confocal laser scanning microscope (Zeiss).

**FDA measurement**

Three days after subcultivation, the *Arabidopsis* cell suspension cultures were diluted 100 times and mixed thoroughly before distribution of 95 µl in 96-well plates. Subsequently, 5 µl of a 1/50 dilution of the bikinin stock solution (1000×) in MSMO medium, DMSO in MSMO medium (used as negative control), or MSMO medium (used as blank control) were added to the cells (1000× final dilution) with a Freedom EVO robot (Tecan). FDA stock solution (2% [w/v] in acetone) was diluted 100 times in target medium and 5 µl was added to 95 µl cell culture. An EnVision 2104 Multilabel Reader (Perkin-Elmer) with the Wallac EnVision manager software package was used to measure fluorescence that was detected with an excitation at 485 nm (band width 14 nm) and emission at 535 nm (band width 25 nm). Relative fluorescence intensities were compared to MSMO (0 min).

**Statistical tests and generation of graphs**

Statistical tests and graphs were generated with Graphpad Prism (version 9.0.1.), except for CETSA MS and phosphoproteomics, of which the statistical tests have been described above. Significant differences were determined with a single-factor analysis of variance (ANOVA) test. The thermal denaturation curves were generated with Boltzmann sigmoid equation with top and bottom constraints set to 1 and 0, respectively, where applicable.

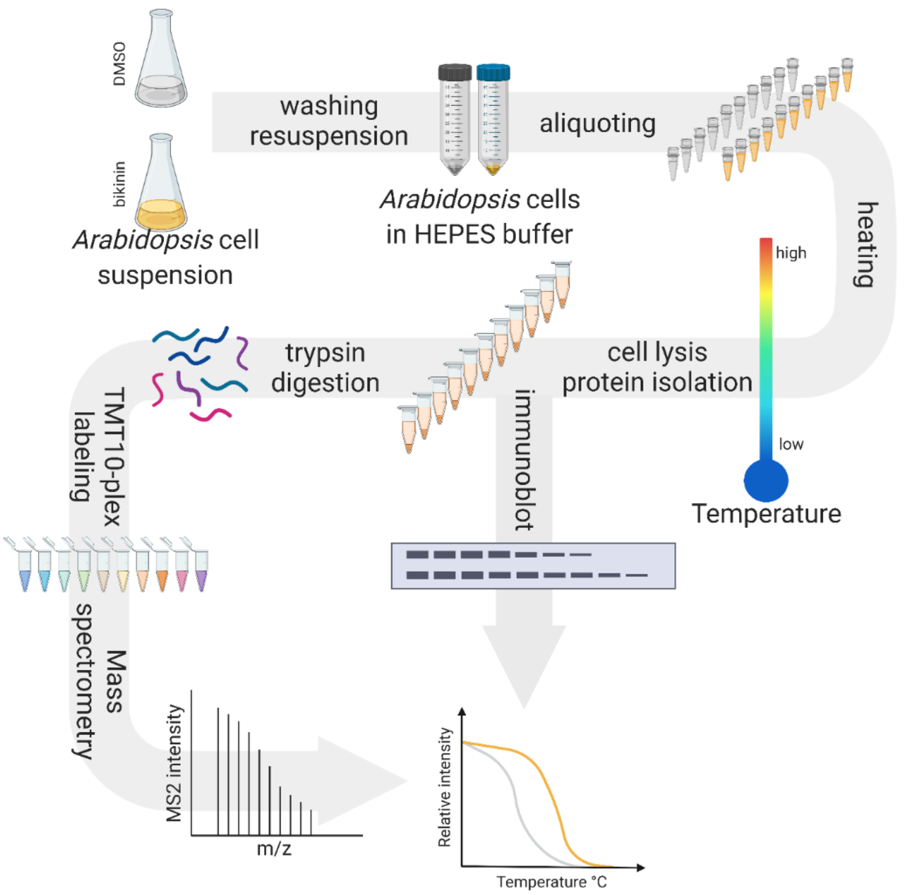


**Fig. S1. Schematic illustration of the cellular thermal shift assay (CETSA) protocol applied to *Arabidopsis* cell suspension cultures.** *Arabidopsis* cell suspension cultures (30 ml) were treated with 50 µM or 250 µM bikinin (as indicated in the text) or 0.1% (v/v) DMSO for 30 min and then washed with protein extraction buffer. For each treatment, cells were divided into 12 or 10 aliquots of 100 µl and heated at 12 and 10 different temperatures for CETSA Western blot (30, 35, 40, 43, 46, 49, 52, 55, 58, 61, 65, and 70°C) and CETSA mass spectrometry (CETSA MS) (25, 30, 35, 40, 45, 50, 55, 60, 70, and 80°C), respectively. Next, the cell samples were subjected to extraction by freezing and thawing. After centrifugation, the supernatants were collected. For CETSA MS, the protein samples were digested with trypsin and labelled with different TMT10 isotope tags for each temperature. Subsequently, all samples from each condition were mixed and analyzed with MS. The obtained reporter ion intensities were used to fit a melting curve and calculate the melting temperature of each protein separately for the two conditions. To generate the melting curves for the CETSA Western blot-detected samples, the proteins amounts were 3 determined by immunoblot with specific antibodies. Figure was created with the BioRender (BioRender.com) software

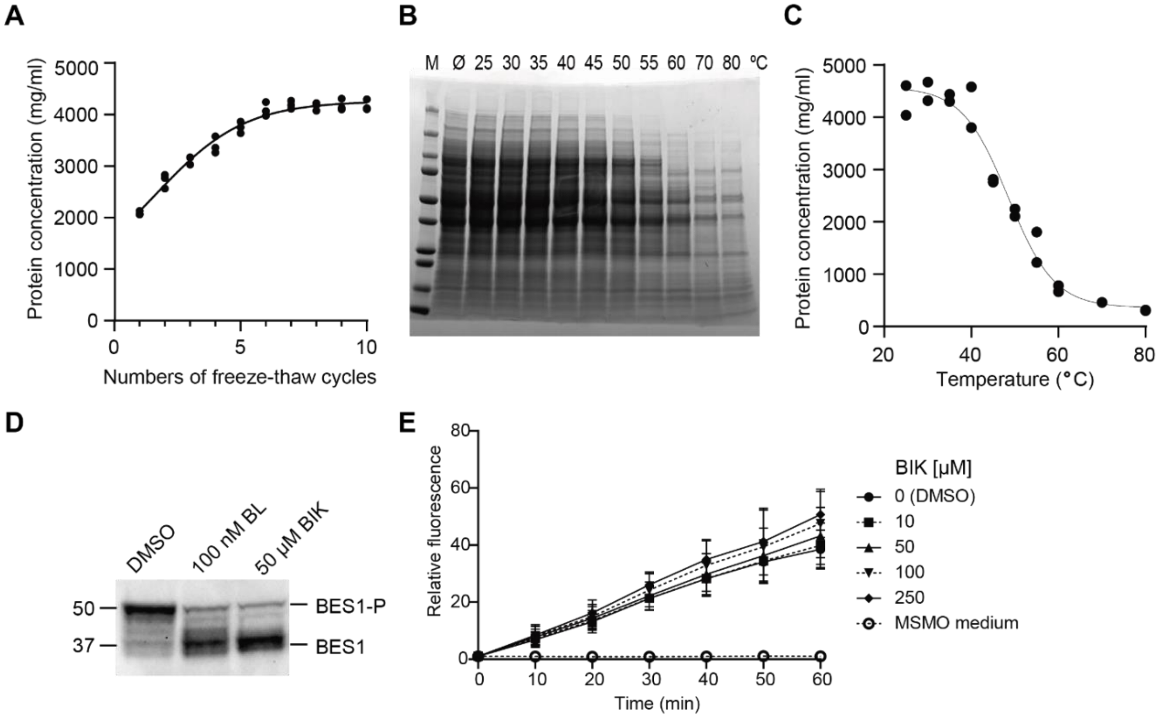


**Fig. S2. Optimization of the cellular thermal shift assay (CETSA) protocol in Arabidopsis cell suspension.** (A) Protein extraction efficiency from Arabidopsis cell suspensions with different freeze-thaw cycles. Individual data points were plotted for three biological replicates. (B) Sodium dodecyl-sulfate polyacrylamide gel electrophoresis (SDS-PAGE) of lysates stained with Coomassie blue. Intact Arabidopsis cells were heated at different temperature for 2 min and lysed by seven freeze-thaw cycles. (C) Quantification of the protein concentrations in (B). Individual data points were plotted for two biological replicates. (D) Accumulation of dephosphorylated BRI1-EMS-SUPPRESSOR1 (BES1) protein as shown by Western blots with anti-BES1 antibodies in Arabidopsis cell suspensions treated with 50 µM bikinin (BIK)- and 100 nM brassinolide (BL) (E) Fluorescein diacetate (FDA) fluorescence measurements of wild type Arabidopsis cell cultures treated with DMSO, different concentrations of bikinin, or the cell culture medium MSMO. Treatment with bikinin did not affect the viability of the Arabidopsis cell suspensions. Values (relatives intensities compared to MSMO [0 min]) are means from three independent experiments. Error bars indicate SD.

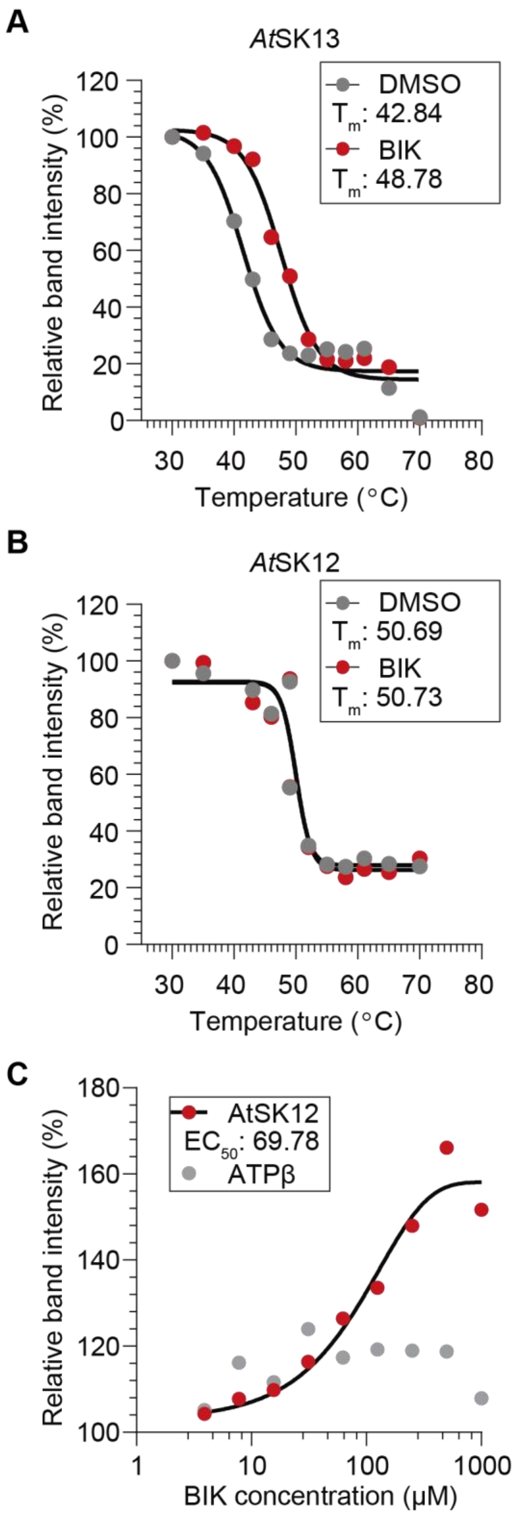


**Fig. S3. Stabilization of AtSK13, but not AtSK12, at low bikinin concentration.** (**A, B**) Thermal denaturation curves for the hemagglutinin (HA)-tagged AtSK13 (A) and AtSK12 (B) both 6 overexpressed in Arabidopsis cell suspension cultures, and for the endogenous ATP synthase subunit β (ATPβ) (A and B) after treatment with 50 µM bikinin (BIK) or 0.1% (v/v) DMSO for 30 min. The relative band intensities from the Western blot analysis were calculated based on the lowest temperature (30°C). The melting temperatures (Tm) are shown on the graphs. (**C**) Thermal denaturation dose-response curves for HA-AtSK12 and the ATPβ generated after treatment with increasing concentrations of BIK for 30 min, the Arabidopsis cell culture were heated at 45°C for 2 min. The relative band intensity from the Western blot analysis was calculated based on the DMSO sample. Tm, melting temperature; EC50, half-maximum response.

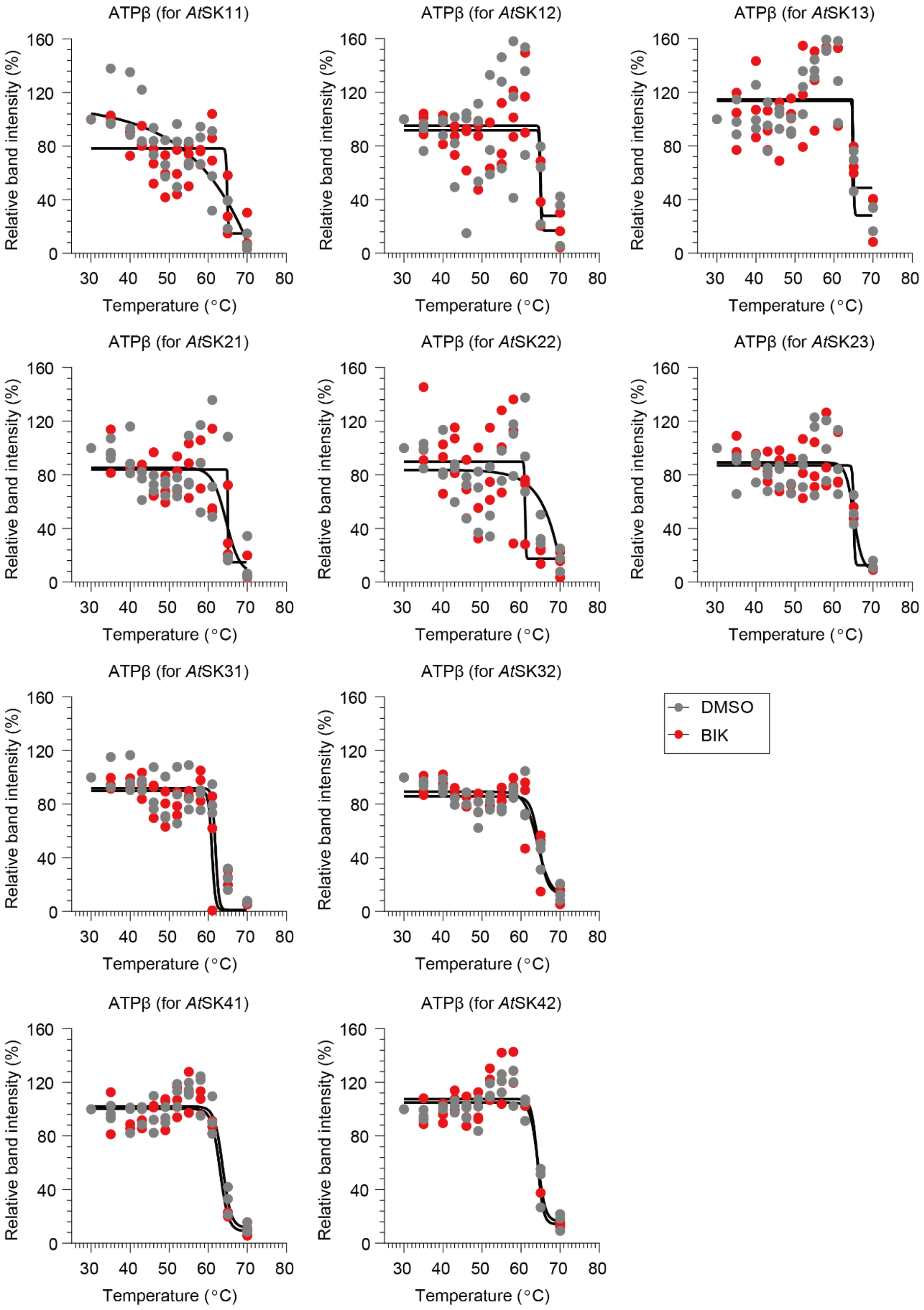


**Fig. S4. Effect of bikinin on the thermal stability of the ATP synthase subunit β.** Thermal denaturation curves for the endogenous ATP synthase subunit β (ATPβ) in the transgenic 8 Arabidopsis cell suspension cultures (as shown in Fig. 1) after treatment with 250 μM bikinin or 0.1% (v/v) DMSO for 30 min. The relative band intensities from the Western blot analysis were calculated based on the lowest temperature (30°C). Individual data points were plotted for three biological replicates.

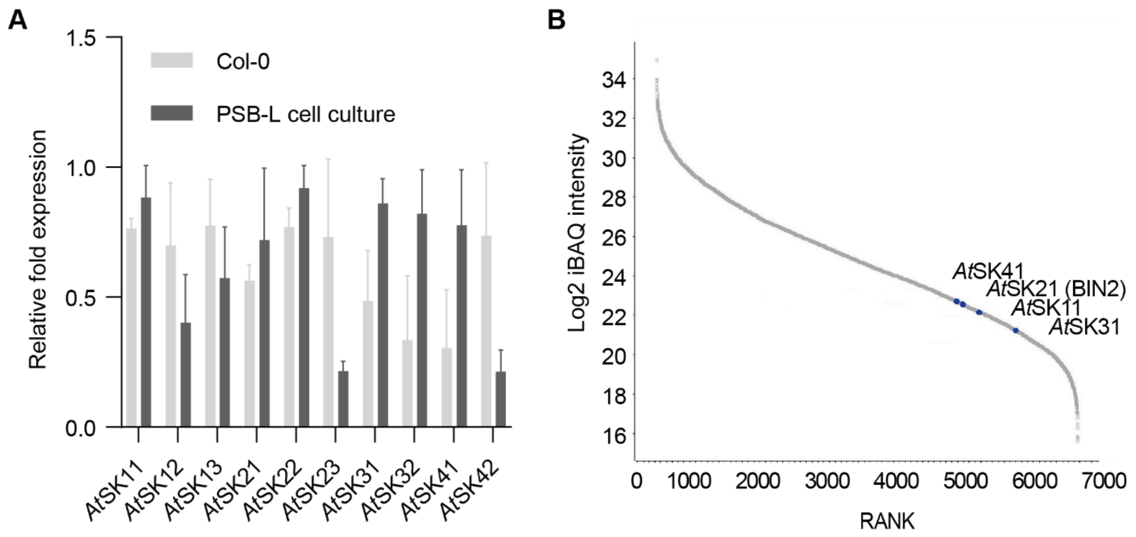


**Fig. S5. Real-Time Quantitative Reverse Transcription (qRT)-PCR and shotgun proteomics analysis of Arabidopsis cell suspension.** (**A**) Transcript levels of the ten AtSKs in Arabidopsis PSB-L cell suspension culture and 5-day-old Arabidopsis (Col-0) seedlings. The transcripts were normalized to the expression of ELONGATION FACTOR 1A (EF1a, AT1G07940). Data represent five biological replicates for each sample. (**B**) Intensity-based absolute quantification (iBAQ) plot representation of all identified proteins in the Arabidopsis cell suspension culture (data file S2). Four of the ten AtSK proteins were identified, albeit at low abundance.

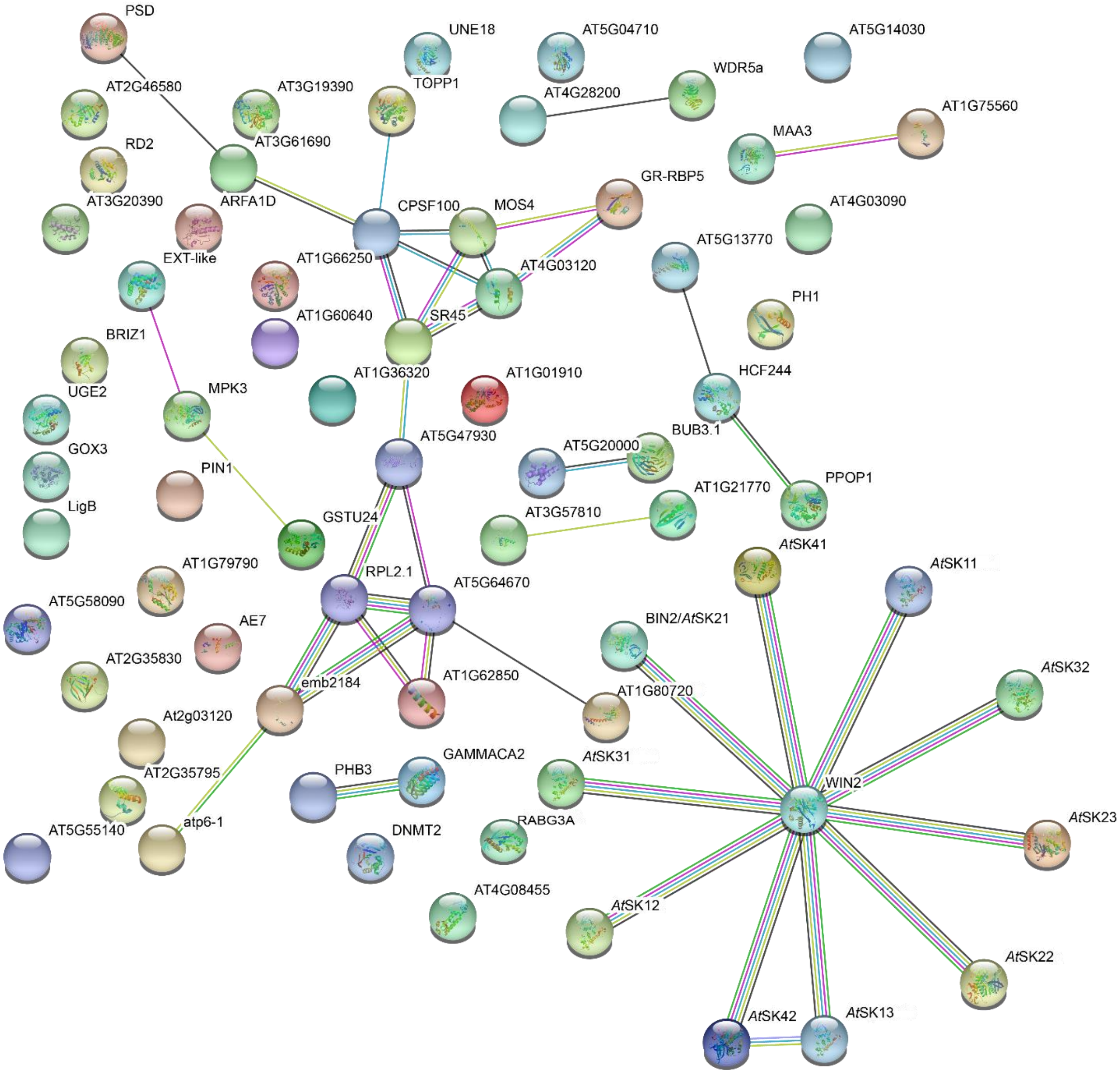


**Fig. S6. STRING analysis of the proteins identified in the cellular thermal shift assay coupled with mass-spectrometry (CETSA MS).** Known and predicted protein-protein interactions according to the STRING database of the 61 proteins identified in the CETSA MS with significant changes in thermal stability (Tm shifts ≥ 2 °C, ANOVA-based F-test $P < 0.01$) (Dataset S1*B*) and the ten AtSK proteins. The analysis was done with STRING (https://version-10-5.string-db.org/).

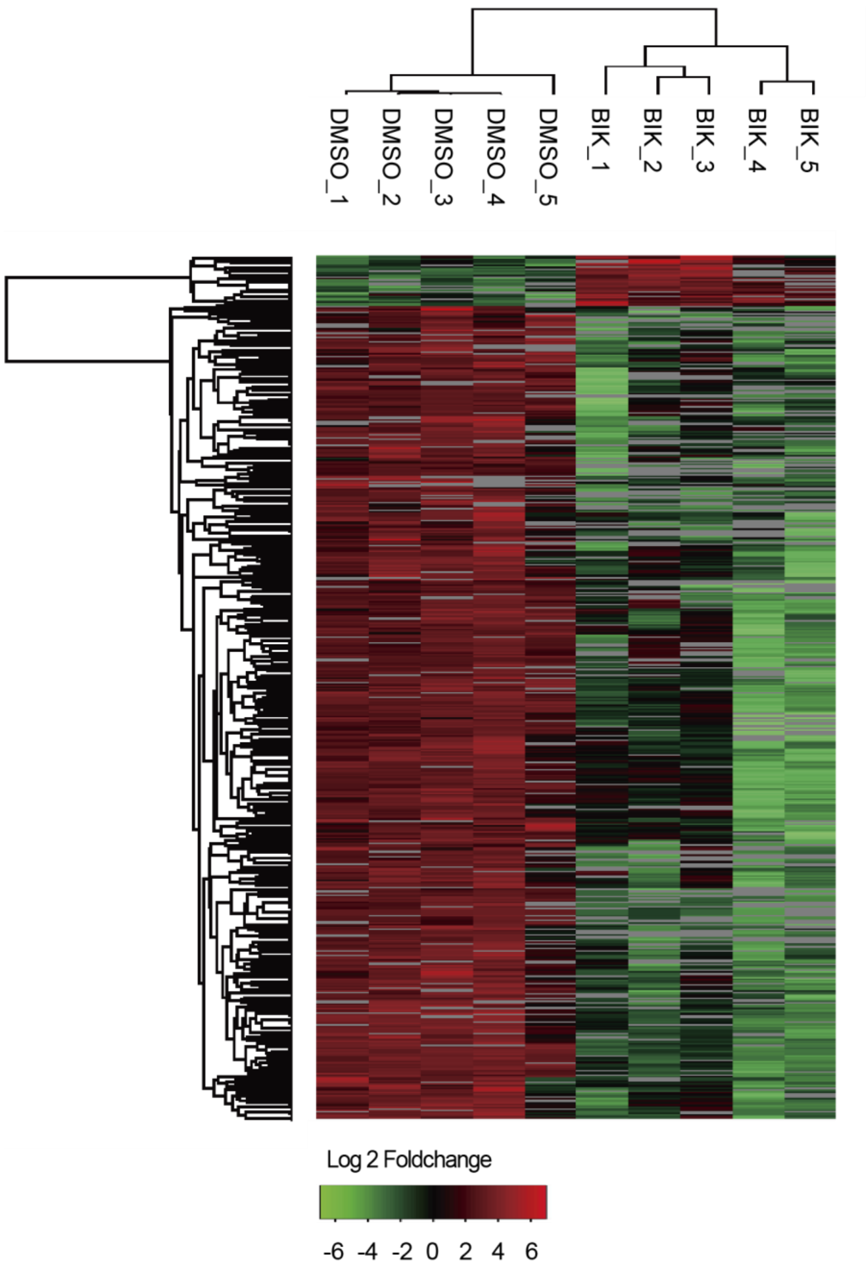


**Fig. S7. Identification of bikinin phosphoproteome.** Hierarchical clustering of 972 downregulated and 101 upregulated phosphorylation sites in Arabidopsis cell suspension cultures treated with 50 µM bikinin (BIK) and compared with DMSO (mock). The columns and rows represent five different biological replicates for each treatment and individual phosphorylation sites, respectively. The scale bar indicates log (base 2)-transformed relative phosphorylation levels of phosphorylation sites. Significant differences were determined with a Student t-test between the treatments, $P < 0.05$.

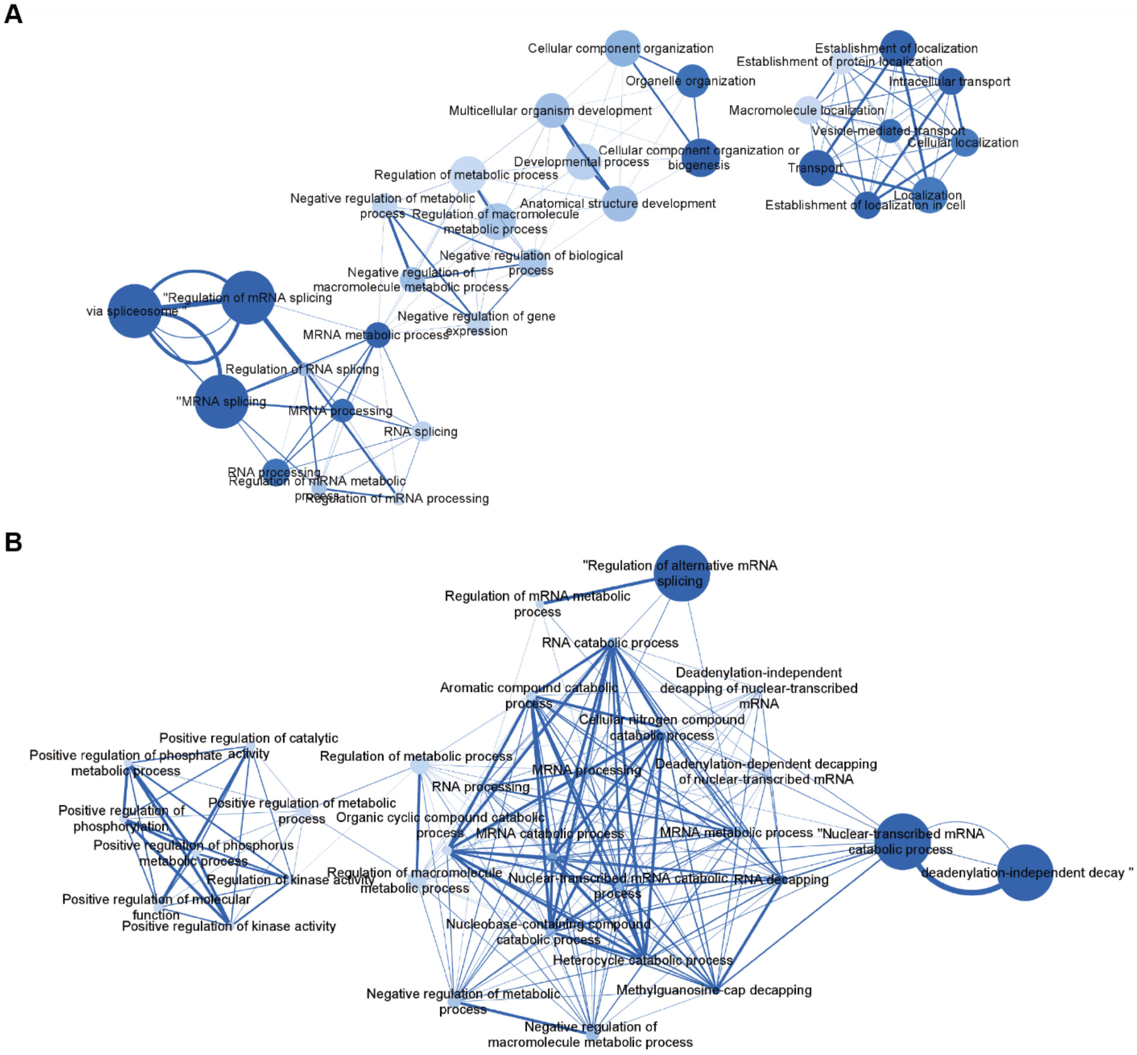


**Fig. S8. Gene ontology (GO) analysis of phosphoproteins up- and downregulated by bikinin.** Network maps of the enriched GO terms for the 665 phosphorylation downregulated proteins (A) and 84 phosphorylation upregulated proteins (B). Only the top 30 significantly enriched terms (FDR adjusted *P* value < 0.05) are shown. Darker nodes indicate more significantly enriched gene sets, diameters correspond with the gene set size and the edge thickness with gene overlapping.

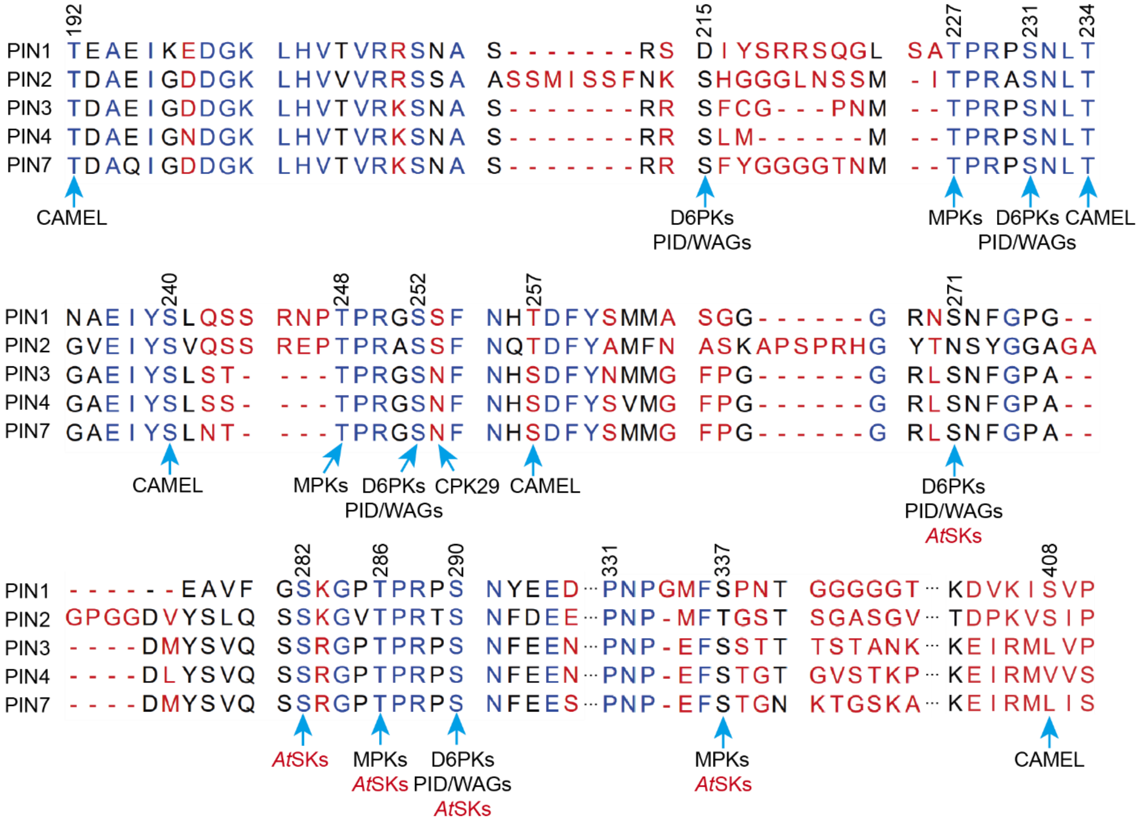


**Fig. S9. Protein sequence alignment of cytoplasmic loops of the long PIN proteins in *Arabidopsis*.** Previously identified phosphorylation sites in the hydrophilic loop of the PIN proteins targeted by the different kinases including AtSKs (this study) as well as PINOID/AGCVIII (PID/WAGs) kinases, D6 Protein Kinases (D6PKs), MITOGEN-ACTIVATED PROTEIN KINASES (MPKs, MPK3/4/6), CANALIZATION-RELATED AUXIN-REGULATED MALECTIN-TYPE RECEPTOR-LIKE KINASE (CAMEL), and CALCIUM-DEPENDENT PROTEIN KINASE 29 (CPK29).

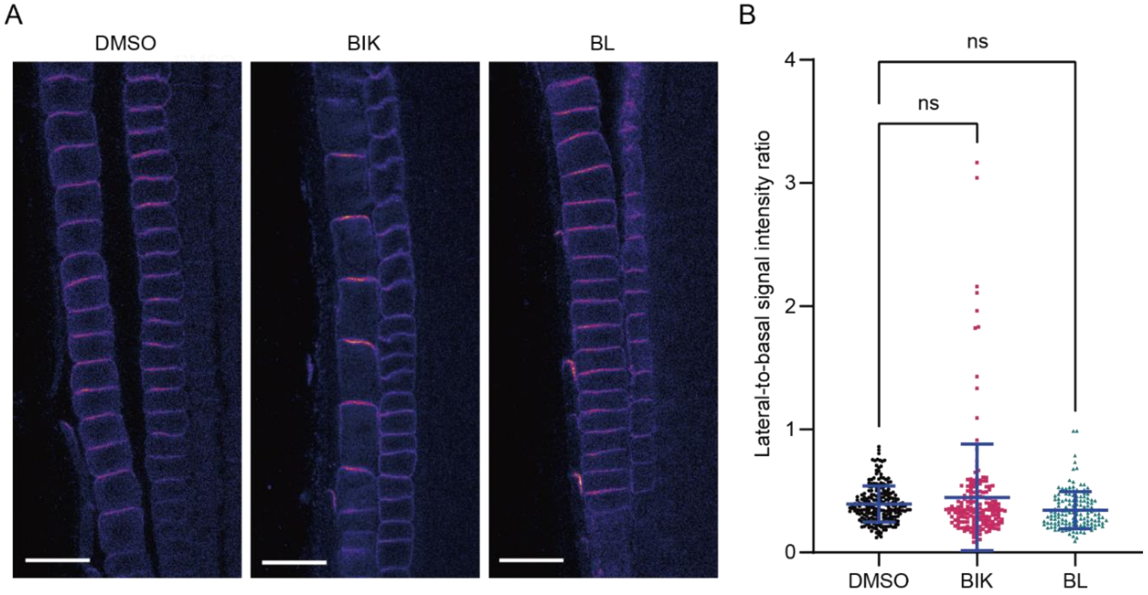


**Fig. S10. Unaffected PIN2 polarity in the Arabidopsis root meristem by brassinolide (BL) and bikinin (BIK) treatments.** (**A**) Immunolocalization of PIN2 in root tips of Arabidopsis after treatment with 50 µM BIK, 10 nM BL, or mock (0.1% [v/v] DMSO) for 12 h. Scale bars, 20 µm. (**B**) Quantification of (A) calculated as the mean ratio of the PIN2 lateral-to-basal signal intensity in the root meristem epidermal cells. Scatter dot plots show all the individual points with the mean and standard errors. One-way ANOVA with Tukey's post hoc test compared to DMSO. *n* > 140 cells corresponding to a minimum of 10 roots per treatment from two independent experiments. ns, not significant.

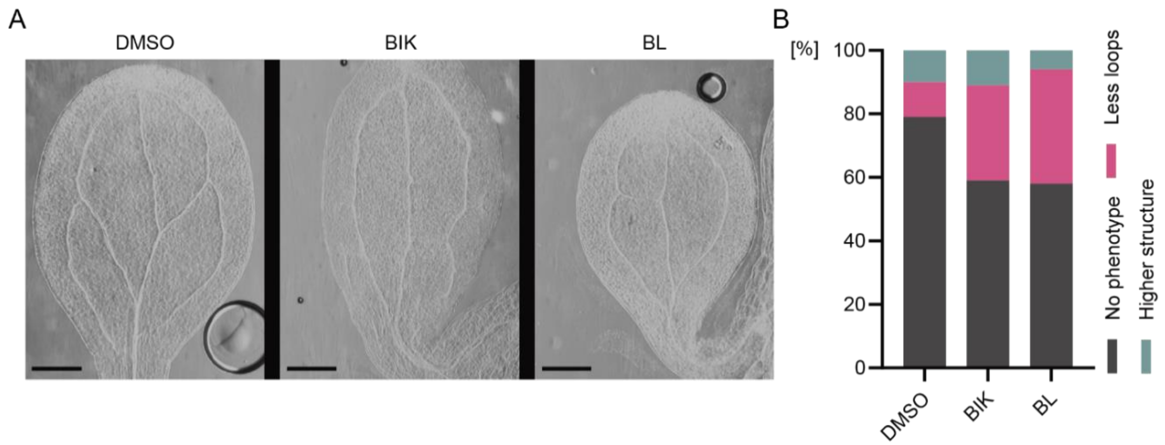


**Fig. S11. Disruption of the vascular bundle patterns in *Arabidopsis* cotyledons by brassinolide (BL) and bikinin (BIK) treatments.** (**A**) Representative images of venation patterning defects in wild-type (Col-0) cotyledons treated with 50 µM BIK, 10 nM BL, or mock (0.1% [v/v] DMSO). Five-day-old Col-0 plants were germinated and grown in liquid medium containing the chemicals for 5 days. Scale bars, 1 mm. (**B**) Quantification of venation defects (n > 40 of each genotype from three independent experiments).

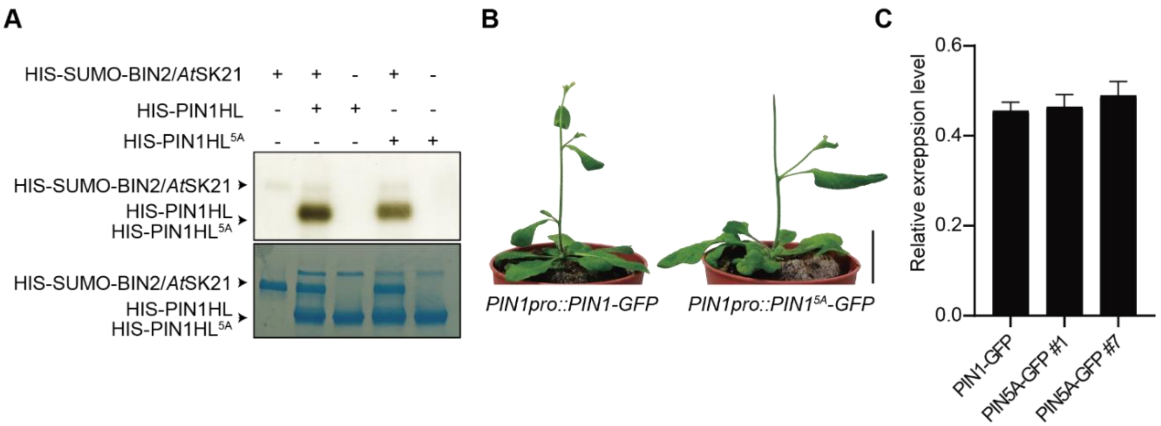


**Fig. S12. Requirement of AtSKs-mediated PIN1 phosphorylation for PIN1 function.** (**A**) Autoradiography (top) and Coomassie brilliant blue staining (bottom) exhibiting reduced phosphorylation of PIN1HL5A by BIN2/AtSK21 in vitro. (**B**) Phenotypes of PIN1pro:: PIN1-GFP and PIN1pro::PIN15A-GFP. Eight of 21 PIN1pro::PIN15A-GFP transgenic plants had naked inflorescence stems in the T1 generation. A photograph of 30-day-old plants is shown. Scale bar, 2 cm. (**C**) PIN1 expression analysis in pPIN1::PIN1-GFP and pPIN1::PIN15A-GFP. Relative transcript levels of the PIN1-GFP and PIN15A -GFP normalized to the ELONGATION FACTOR 1A (EF1a, AT1G07940) expression in 5-day-old seedlings of pPIN1::PIN1-GFP and pPIN1::PIN15A-GFP, respectively. Values are means from three independent experiments. Error bars indicate SD.

**Table S1. Primers used in this study**

| primers | Sequence |
|---|---|
| **Gibson-AtSK11-F** | ATTGAGGCTCACAGAGAACAGATTGGTGGATCCATGGCGTCAGTGGGTATAGCTC |
| **Gibson-AtSK11-R** | TACTTTCTGTTCGACTTAAGCATTATGCGGCCGCTCACAAACCGAGCCAAGGACAC |
| **Gibson-AtSK12-F** | ATTGAGGCTCACAGAGAACAGATTGGTGGATCCATGGCCTCGGTGGGCATAGAGC |
| **Gibson-AtSK12-R** | TACTTTCTGTTCGACTTAAGCATTATGCGGCCGCTCACAAACTGAGCCACGGACAT |
| **Gibson-AtSK13-F** | ATTGAGGCTCACAGAGAACAGATTGGTGGATCCATGGCTTCTGTGGGAACATTACC |
| **Gibson-AtSK13-R** | TACTTTCTGTTCGACTTAAGCATTATGCGGCCGCTTAGAGAGCGAGGAAGGAACATTG |
| **Gibson-AtSK21-F** | ATTGAGGCTCACAGAGAACAGATTGGTGGATCCATGGCTGATGATAAGGAGATGC |
| **Gibson-AtSK21-R** | TACTTTCTGTTCGACTTAAGCATTATGCGGCCGCTTAAGTTCCAGATTGATTCAAGA |
| **Gibson-AtSK22-F** | ATTGAGGCTCACAGAGAACAGATTGGTGGATCCATGGCCTCATTACCATTGGG |
| **Gibson-AtSK22-R** | TACTTTCTGTTCGACTTAAGCATTATGCGGCCGCTTAACTGTTTTGTAATCCTGTG |
| **Gibson-AtSK23-F** | ATTGAGGCTCACAGAGAACAGATTGGTGGATCCATGACTTCGATACCATTGGG |
| **Gibson-AtSK23-R** | TACTTTCTGTTCGACTTAAGCATTATGCGGCCGCCTAGGGTCCAGCTTGAAATGGA |
| **Gibson-AtSK31-F** | ATTGAGGCTCACAGAGAACAGATTGGTGGATCCATGAATGTGGTGCGGAGATTAAC |
| **Gibson-AtSK31-R** | TACTTTCTGTTCGACTTAAGCATTATGCGGCCGCTCATTTCCTTGCATGCTCAGG |
| **Gibson-AtSK32-F** | ATTGAGGCTCACAGAGAACAGATTGGTGGATCCATGAACGTGATGCGTCGTCTC |
| **Gibson-AtSK32-R** | TACTTTCTGTTCGACTTAAGCATTATGCGGCCGCCTAAGAGCTACTTCCCGTTCC |
| **Gibson-AtSK41-F** | ATTGAGGCTCACAGAGAACAGATTGGTGGATCCATGGCATCCTCTGGACTGGGA |
| **Gibson-AtSK41-R** | TACTTTCTGTTCGACTTAAGCATTATGCGGCCGCTTACGAATGCAAAGCCATGAAG |
| **Gibson-AtSK42-F** | ATTGAGGCTCACAGAGAACAGATTGGTGGATCCATGGAATCTCATCTGGGAAATG |
| **Gibson-AtSK42-R** | TACTTTCTGTTCGACTTAAGCATTATGCGGCCGCTTACGAGTGTAATGCCATGAAG |
| **WIN2-ATTB1** | GGGGACAAGTTTGTACAAAAAAGCAGGCTTAATGGGATATCTGAATTCTGTTTTG |
| **WIN2-ATTB2** | GGGGACCACTTTGTACAAGAAAGCTGGGTAGGTTGATGAGTCACCGGAGA |
| **AtSK11-qPCR-fwd** | GCGTCAGTGGGTATAGCTCC |
| **AtSK11-qPCR-rev** | ACAACACGCTCAGCCATGTA |
| **AtSK12-qPCR-fwd** | TCCGTTGTGCTGCTCTTGAT |
| **AtSK12-qPCR-rev** | AATCGCGCATTCGGATCTCT |
| **AtSK13-qPCR-fwd** | AAGGCGAGCCAAACATCTCA |
| **AtSK13-qPCR-rev** | GAGGCTGTCCCAGAAGCAAT |
| **AtSK21-qPCR-fwd** | CACAAAAGGATGCCCCCAGA |
| **AtSK21-qPCR-rev** | TTGAAGAGAGGCGGGAAAGG |
| **AtSK22-qPCR-fwd** | GACCTTGCATCTCGGCTTCT |
| **AtSK22-qPCR-rev** | GCTCCATTGAAGCTCCACCT |
| **AtSK23-qPCR-fwd** | AACTCGCGAAGAAATCCGGT |
| **AtSK23-qPCR-rev** | AACCTTATGCCAAGGGTGGG |
| **AtSK32-qPCR-fwd** | AGAGACGAGCGAAATGCCAA |
| **AtSK32-qPCR-rev** | CGCTGGGCCATGTATGAGAT |
| **AtSK31-qPCR-fwd** | AGAGACCCGAGAGCATCCTT |
| **AtSK31-qPCR-rev** | GACGCAGTTCAACAGATGCC |
| **AtSK41-qPCR-fwd** | ACTGGGAAATGGAGTAGGCAC |
| **AtSK41-qPCR-rev** | CCCTTATCCTCGTCTCAGCC |
| **AtSK42-qPCR-fwd** | GCCACTTCCTCCGCTATTCA |
| **AtSK42-qPCR-rev** | GGACAAGTCGGTCCACAGTT |
| **EF1a-qPCR-fwd** | TGAGCACGCTCTTCTTGCTTTCA |
| **EF1a-qPCR-rev** | GGTGGTGGCATCCATCTTGTTACA |
| **PIN1-S270A-F** | CGGAACGCTAACTTTGGTCCTGGAGAAGC |
| **PIN1-S270A-R** | CAAAGTTAGCGTTCCGACCACCACCAGAAG |
| **PIN1-3A-F** | GCTAAAGGTCCTGCTCCGAGACCTGCCAACTACGAAGAAGACGGTGG |
| **PIN1-3A-R** | GGCAGGTCTCGGAGCAGGACCTTTAGCACCAAACACAGCTTCTCCAGGA |
| **PIN1-S337A-F** | ATGTTTGCGCCCAACACTGGCGGTGGTGGA |
| **PIN1-S337A-R** | AGTGTTGGGCGCAAACATCCCTGGGTTCGGCG |
